# Cyclo-stationary distributions of mRNA and Protein counts for random cell division times

**DOI:** 10.1101/2025.06.06.658238

**Authors:** Syed Yunus Ali, Aditya Saran, Ashok Prasad, Abhyudai Singh, Dibyendu Das

**Affiliations:** Department of Physics, Indian Institute of Technology Bombay, Powai, Mumbai 400076, India; School of Biomedical and Chemical Engineering, Colorado State University, Fort Collins, Colorado 80521, USA; Department of Electrical and Computer Engineering, University of Delaware, Newark, DE 19716, USA; Max Planck Institute for the Physics of Complex Systems, 01187 Dresden, Germany

## Abstract

There is a long history of using experimental and computational approaches to study noise in single-cell levels of mRNA and proteins. The noise originates from a myriad of factors: intrinsic processes of gene expression, partitioning errors during division, and extrinsic effects, such as, random cell-cycle times. Although theoretical methods are well developed to analytically understand full statistics of copy numbers for fixed or Erlang distributed cell cycle times, the general problem of random division times is still open. For any random (but uncorrelated) division time distribution, we present a method to address this challenging problem and obtain exact series representations of the copy number distributions in the cyclo-stationary state. We provide explicit cell age-specific and age-averaged results, and analyze the relative contribution to noise from intrinsic and extrinsic sources. Our analytical approach will aid the analysis of single-cell expression data and help in disentangling the impact of variability in division times.

## I. Introduction

Advances in technologies of single-cell RNA sequencing and single-molecule fluorescence in situ hybridization to quantify mRNA levels and fluorescent proteomic imaging, mass cytometry and mass spectrometry to quantify protein levels in individual cells, have unmasked tremendous intercellular variability within isogenic populations over the last two decades ***Raj et al. (2008); Raj and Van Oudenaarden (2008); Lovatt et al. (2014); Cao et al. (2017); Mahdessian et al. (2021); Suen et al. (2021); Deloupy et al. (2020***). Understanding the different sources of stochasticity that drive this variability is key to the analysis of single-cell transcriptomic and proteomics data, and using stochasticity as a tool to infer complex regulatory networks ***Munsky et al. (2012); Padovan-Merhar and Raj (2013); Saint-Antoine and Singh (2020***). Stochastic expression has been implicated in diverse emerging medical problems, such as, cancer drug resistance ***Shaffer et al. (2017); Farquhar et al. (2019); El Meouche et al. (2024); Chang et al. (2022***), microbial persistence and replication of human viruses ***El Meouche et al. (2016); Singh and Weinberger (2009***). Advancing analytical tools for understanding and modeling these inherent noise mechanisms can directly impact controlling cell-to-cell variation for therapeutic benefit.

The protein and mRNA copy numbers in cells are determined by a series of coupled stochastic chemical processes, leading to the above mentioned significant cell-to-cell variability. The transcriptional and translational noise arise due to multiple factors – genes switching between transcriptionally active and inactive states, rapid decay of short-lived mRNA leaving behind long-lived proteins making them appear in bursts, and other factors like RNA splicing and post-transcriptional regulation by micro-RNAs ***Swain et al. (2002); Raser and O’Shea (2004); Raj et al. (2006); Cai et al. (2006); Taniguchi et al. (2010); Sanchez and Golding (2013); Storz et al. (2004***). Theoretical studies of simple models (ignoring certain complexities) have obtained analytical moments and probability distributions of mRNA and protein count in the steady state ***Thattai and Van Oudenaarden (2001); Paulsson (2005); Sanchez et al. (2011); Shahrezaei and Swain (2008); Bokes et al. (2012); Raj et al. (2006); Singh et al. (2013***), as well as for transient perturbations ***Singh et al. (2012***) and along the cell cycle ***Wang et al. (2023***).

In addition to intrinsic noise in gene expression specific to every gene, there are extrinsic factors affecting all genes. Two significant contributors to cell-to-cell variability in copy number, on which we focus in this paper, are noise incorporated through variable cell-cycle times ***Hawkins et al. (2009); Tsukanov et al. (2011); Reshes et al. (2008a); Roeder et al. (2010); Stukalin et al. (2013); Yates et al. (2017); Thomas (2019***), and random partitioning of copy numbers to daughter cells after cell division ***Huh and Paulsson (2011a***,b); ***Zopf et al. (2013); Cookson et al. (2009***). The noise associated with cell division has also been considered by various theoretical studies on copy number fluctuations ***Berg (1978); Rigney (1979); Bertaux et al. (2018); Dessalles et al. (2020); Soltani and Singh (2016); Beentjes et al. (2020); Cao and Grima (2020); Jia et al. (2022); Wang et al. (2023***), discussed in more detail below. In this paper, we revisit this problem.

Within a cell cycle with a finite division time, steady states are not attained for mRNA and protein counts. Yet after many successive cycles of division, a *cyclo-stationary* state is reached when time independent distributions are attained for every cell ‘age’ – the age is zero at birth and maximum just before division. The statistics of copy numbers in the cyclo-stationary state have been of central interest in the literature mentioned above.

What decides the instant of cell division still remains an intriguing question. Cell division has been argued to be triggered by a ‘timer’, a ‘sizer’, or an ‘adder’ mechanism. In the ‘timer’ scenario, it is assumed that cell division happens after fixed times *T* ***Fantes and Nurse (1977***). This is a common assumption and has been extensively used in the theoretical literature ***Berg (1978); Rigney (1979); Huh and Paulsson (2011a); Beentjes et al. (2020); Cao and Grima (2020); Wang et al. (2023); Johnston and Jones (2015***). Yet quite generally, cell division times are known to be random, and also dependent on the cell size to ensure size homeostasis ***Männik et al. (2024); Vargas-Garcia et al. (2018); Tsukanov et al. (2011); Reshes et al. (2008a***,b); ***Roeder et al. (2010); Zilman et al. (2010); Hawkins et al. (2009); Stukalin et al. (2013); Liu et al. (2024); Nieto et al. (2024b***,b). In the ‘sizer’ view, division occurs when a cell size threshold is crossed ***Campos et al. (2014); Fantes and Nurse (1977***), while in the ‘adder’ view, it happens when the additional cell size growth from the initial birth size crosses a threshold ***Cadart et al. (2018); Taheri-Araghi et al. (2015); Nieto et al. (2025***). In the literature, ‘threshold crossing’ has been modeled in different ways. One approach splits the cell cycle into a sequence of *N* steps with exponentially distributed time intervals, leading to cell division and copy number partitioning at the end of the *N*-th step, which therefore acts like a threshold. The *N* steps are not actual cell cycle phases, but a model assumption. The total cell division time follows an Erlang or Hypoexponential distribution in such cases ***Yates et al. (2017); Soltani and Singh (2016); Soltani et al. (2016); Perez-Carrasco et al. (2020); Beentjes et al. (2020); Jia and Grima (2021); Jia et al. (2022); Nieto et al. (2024a***). In contrast, another view treats cell division as a *first-passage time problem* ***Gardiner (1985); Ghusinga et al. (2016***) (i.e., first threshold crossing of key regulatory protein). For an auto-catalytic growth process – this leads to a Beta-exponential distribution of division times ***Iyer-Biswas et al. (2014a***,b). Some recent works have assumed stochastically fluctuating thresholds instead of a fixed threshold, and proposed some empirically relevant distributions of cell cycle times ***Luo et al. (2023); Biswas and Brenner (2024***).

The variability of cell cycle times is thus a fact howsoever diverse may be the cause, yet theoretical works in the past have not treated this aspect in full generality. One study considered random cell division times and obtained exact moments of the copy numbers, but nevertheless assumed deterministic growth kinetics and deterministic partitioning ***Antunes and Singh (2015***). A large body of analytical work has completely ignored the randomness in cell cycle times. The coefficient of variation was studied in ***Huh and Paulsson (2011a***) for fixed division times, comparing the relative role of gene expression noise versus binomial partitioning. Under the same assumption of fixed times, the generating functions for the distributions of mRNA and proteins in the cyclo-stationary state were derived for various models ***Berg (1978); Rigney (1979); Beentjes et al. (2020); Cao and Grima (2020); Jędrak et al. (2019); Wang et al. (2023***). Specifically, constitutive bursty protein production ***Beentjes et al. (2020***), a two-stage model for mRNA synthesis with active and inactive transcription states ***Cao and Grima (2020***), and a three-stage model for protein synthesis ***Wang et al. (2023***), were studied. The exact generating functions were derived for age-specific and age-averaged cases, and in the presence or absence of gene duplication. The desired probability distributions of copy number were then obtained through numerical derivatives of the generating functions.

Random cell cycle times were treated in another set of works, but for a special class of distributions – Erlang, mixed Erlang, and Hypoexponential. These distributions permit an alternate representation of the cell cycle by a Markov chain of *N* stages each with an exponentially distributed lifetime. This technical simplicity, along with further assumption of steady state in each stage, led to exact moments and cumulants ***Soltani and Singh (2016); Soltani et al. (2016); Perez-Carrasco et al. (2020***). Under the same assumptions, the generating function for ‘age-averaged’ protein distributions, for bursty synthesis without degradation, were derived in ***Beentjes et al. (2020***). The complexities of volume-dependent gene expression, gene duplication, and dosage compensation were also incorporated in two of these papers ***Jia et al. (2022); Beentjes et al. (2020***). The power spectra of the copy number auto-correlation function in the cyclo-stationary state was studied in ***Jia and Grima (2021***).

Although the literature discussed above has made valuable contributions to our theoretical understanding of the problem and compared with some experimental data, it is evident that an analytical approach is lacking to treat arbitrary random division times which may arise in experiments. Even the only case widely studied, namely Erlang (and Hypoexponential), used an assumption of steady-state for each of the constitutive stages of the process, and thereby was limited to obtaining the ‘age-averaged’ distributions of the copy number. Cell age-specific distributions are preferable as age-averaged ones may be derived from those, but the other way around is not possible. Given this, and the fact that other empirically relevant distributions have been reported ***Iyer-Biswas et al. (2014a); Biswas and Brenner (2024); Luo et al. (2023***), there is sufficient motivation to take a fresh look at the problem in this paper.

We adapt the framework of generating functions used for solving the cyclo-stationary distributions of copy numbers to the case of random division times, and firstly indicate why going beyond the case of fixed cell cycle times has remained technically challenging. We then develop a method to tackle this problem mathematically and evaluate the cyclo-stationary distributions as certain *analytically exact series* (Sec. II). Instead of enumerating the derivatives of the generating functions ***Beentjes et al. (2020); Cao and Grima (2020); Wang et al. (2023***), the method now requires summing the relevant series directly. It works for any random (uncorrelated) cell cycle time distribution – the Erlang distribution can now be plugged in directly without splitting it into steps, just like Beta exponential, Lognormal, or other empirically relevant distributions (Sec. III, VI D) as we show in the paper. We also show that for fixed division times, the cyclo-stationary mRNA distribution is exactly Poisson (Sec. IV). We provide explicit results for the basic gene expression model of mRNA synthesis and bursty protein production, with both having finite degradation rates ***Friedman et al. (2006); Shahrezaei and Swain (2008***) (Sec. IV, V). The cyclo-stationary distributions we obtain are age-specific – at cell birth, before division, or any time in between (Sec. VI A). We show that the age-averaged results may be obtained using suitable age frequency functions (Sec. VI B). Exact noise (*CV* ^2^) formulas and separate contributions to it from intrinsic and extrinsic sources are calculated – the part due to random cell cycle times depending on its distribution can be as high as 50% (Sec. VI C)). The Skewness shows a non-monotonicity with mean division times, which is more pronounced with higher variability of cell cycle times – thus can serve as a diagnostic of the variability itself (Sec. V). The impact of correlations in successive cycle times is studied in Sec. VI E.

## II. The method to treat random cell cycle times, & cyclo-stationary distributions at cell Birth and before division

In this section, we present a general framework within which cyclo-stationary distributions of mRNAs or proteins may be analytically derived, for *random* cell cycle times. Let us denote the integer copy number of either mRNA or protein by *y*(*t*). Later we will replace *y*(*t*) = *m*(*t*) for mRNA and *y*(*t*) = *n*(*t*) for protein. As shown schematically in Fig. 1, *y*(*t*) grows from a value *y*_+,*i*−1_ at the beginning of the *i*^*th*^ cell cycle to a value of *y*_−,*i*_ at its end. Then as the cell divides, *y*_−,*i*_ binomially partitions between two daughter cells with copy numbers *y*_+,*i*_ and *x*_+,*i*_ = *y*_−,*i*_ − *y*_+,*i*_ ***Golding et al. (2005***). This process repeats over several generations through repeated cell divisions as depicted in Fig. 1, i.e. *i* = 1, 2, … All the details of gene expression within a cell cycle through transcription and translation are described by the probability distribution 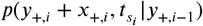. Here 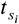 denotes the duration of the *i*^*th*^ cell cycle. Using the binomial distribution 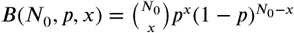 with *p* = 1/2, we may relate the distributions of copy numbers 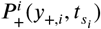 and 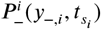 just after and just before the *i*^*th*^ cell division respectively, to the distribution 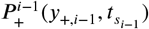 after the (*i* − 1)^*th*^ cycle as follows:

**Figure 1.**
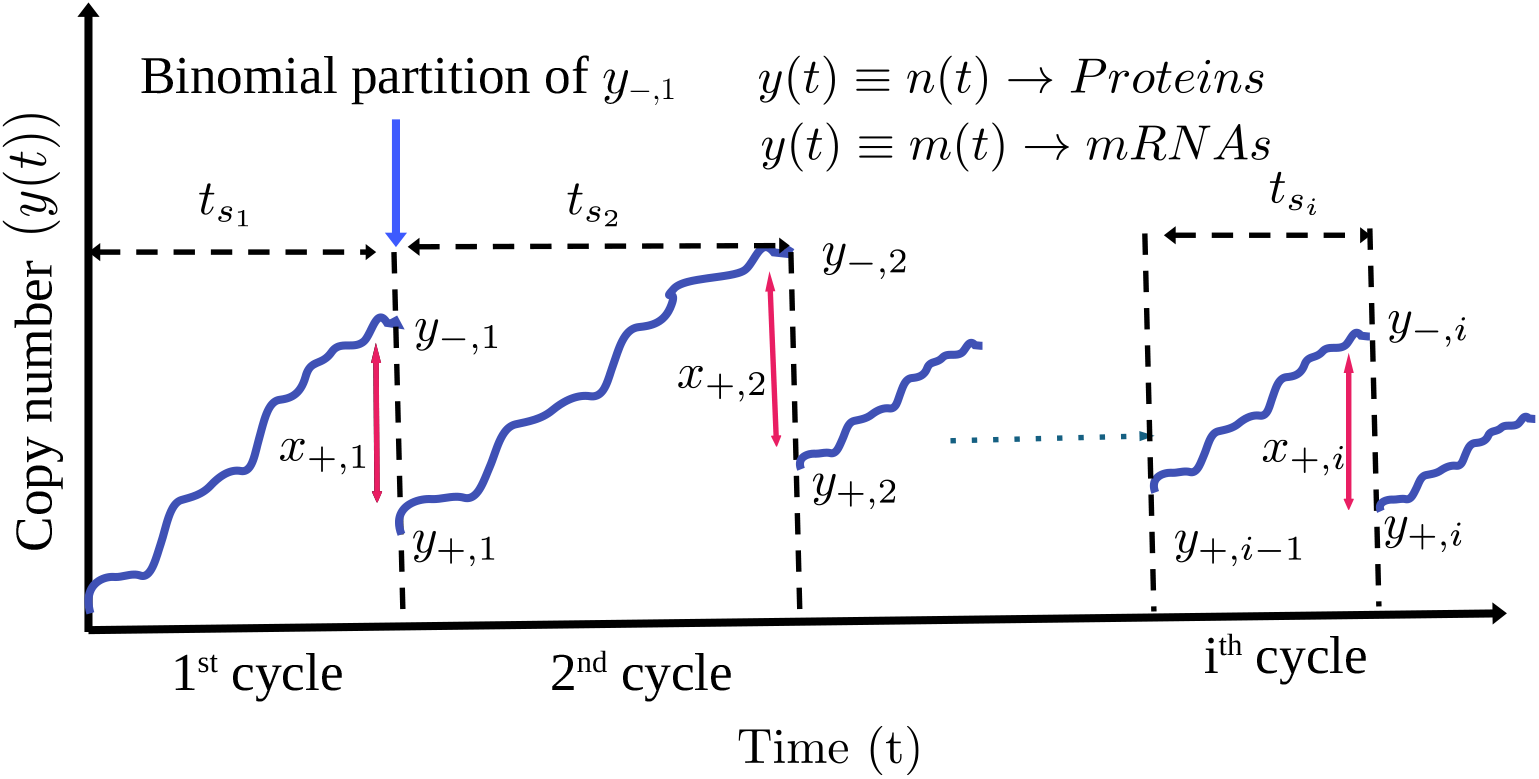
Schematic diagram of time evolving copy number, of either mRNA (*m*(*t*)) or proteins (*n*(*t*)) in successive generations, interrupted by cell-divisions when the number *y*_−,*i*_ binomially divides to *x*_+.*i*_ and *y*_+,*i*_ in the two daughter cells. The durations 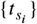 of the cell cycles are random, and drawn from a probability distribution *g*(*t*_*s*_). After several generations, i.e. *i* ≫ 1, the copy numbers attained cyclo-stationary distributions 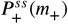 and 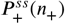, which are studied in the text.

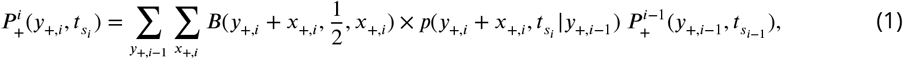

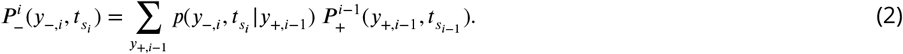

Cyclo-stationary state is attained for *i* ≫ 1, when the above two distributions approach steady (cycle independent) forms 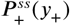 (for new born cells) and *P* ^*ss*^(*y*_−_) (for most mature cells before division). Our aim in this section is to solve for these.

To reemphasize, there are three sources of stochasticity – gene expression controlling the evolution of *y*(*t*) within every cycle, the random binomial partitioning at every division step, and the random cell cycle times 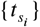. Let the distribution of these division times be 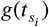– identical for every cycle *i*. As noted above, various works ***Berg (1978); Rigney (1979); Beentjes et al. (2020); Cao and Grima (2020); Wang et al. (2023***) studied the special case of fixed cell cycle times 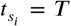, i.e. 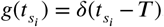. In that case, generating function for the probability of copy number in the *i*^th^ cycle could be recursively related to that of the (*i* − 1)^th^ cycle. We will show below how such recursive proportionality is not realized for general random 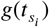.

Next we highlight an assumption, implicitly made in almost all past literature, that the successive cell division times are uncorrelated, i.e. the two-point function 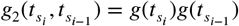. In such cases, starting from Eqs. 1 and 2, it may be shown (see supplementary material (SM) Sec.-IA ***sup (2025***)) that the cyclo-stationary copy number distributions 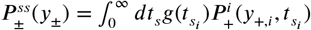 satisfy:

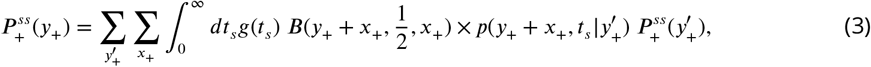

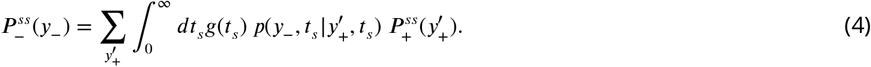

Here subscripts *i* has been dropped by setting 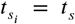 and *y*_±,*i*_ = *y*_±_ to indicate the history independence. Next we need information of the model of gene expression. For a concrete study, we suppose the evolving copy number distribution 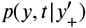 has a generating function 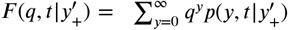 of the form

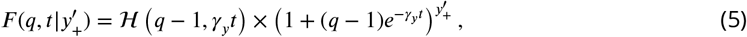

which is indeed the case for gene expression models of mRNA and protein, studied below. The function ℋ (.) is specific to the process. Given Eq. 5, it may then be shown (SM Sec.-IB ***sup (2025***)) that the generating functions 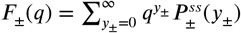 of the cyclo-stationary distributions follow:

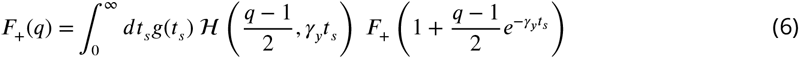

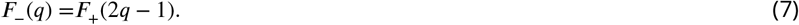

For fixed cell cycle time *T* i.e. *g*(*t*_*s*_) = *δ*(*t*_*s*_ − *T*), Eq. 6 solves exactly for the generating function (see SM Sec-IC)

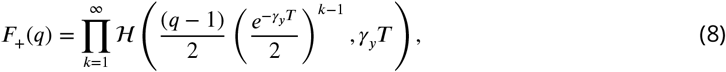

reminiscent of earlier works ***Berg (1978); Beentjes et al. (2020); Cao and Grima (2020***). Consequently, the steady-state probability is obtained through its derivatives: 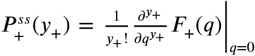.

In contrast, for a random *t*_s_, with a general distribution *g*(*t*_s_), the *F*_+_ in the right-hand side of the Eq. 6 on repeated iteration leads to nested integrals as shown in Eq. 21 of SM Sec-IC ***sup (2025***), which are generally intractable. Thus the function *F*_+_(*q*) in general seems hard to find analytically.

We bring in a useful insight to this challenging problem by noting that the mathematical structure of the problem at hand is very similar to the one arising in the process of synaptic vesicle fusion and release across chemical synapses, on cyclic stimulation by action potentials ***Bird et al. (2016); Krächan et al. (2017); Rijal et al. (2024); Vahdat et al. (2025***). The size of the ready-release pool of synaptic vesicles in the pre-synaptic neuron evolves as the copy number in Fig. 1. The arrival of an action potential at the pre-synaptic terminal causes sudden vesicle release and reduction of the pool size, just like the reduction in copy number by partitioning during cell division. The number of vesicles fused and released are binomially distributed, resembling the binomial partitioning of the copy number to daughter cells. The interspike intervals between arrival of action potentials are like random cell cycle times. In the study of statistics of the quantal content of synaptic vesicle release, a similar equation as Eq. 6 arises ***Rijal et al. (2024***), and we follow an idea found useful in that context.

Note that in Eq. 6, the argument *q* of the generating function *F*_+_ on the left side, maps to another argument 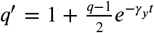 on the right side. A fixed point of this map is *q*^′^ = *q* = 1. Hence, a useful way to proceed analytically is to do an alternate series expansion of *F*_+_(*q*) about *q* = 1 as follows:

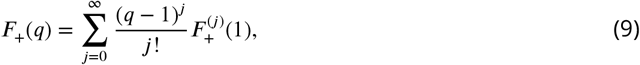

Substituting the above on both sides of Eq. 6, an exact recursion relation between the coefficients 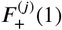 of the following general form may be obtained (the algebra of which will be demonstrated through examples below):

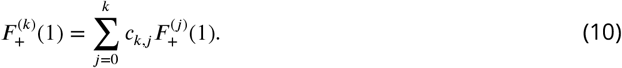

The information of the distribution *g*(*t*_*s*_) gets embedded in the coefficients *c*_*k,j*_. Thus, the key to solving the cyclo-stationary distributions of the copy number for random cell cycle times is to evaluate the coefficients 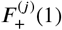 from the above equation Eq. 10. Using those, as shown in SM Sec-ID ***sup (2025***), the final distributions are given by:

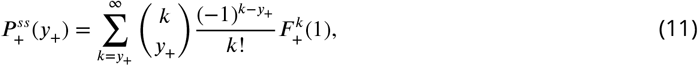

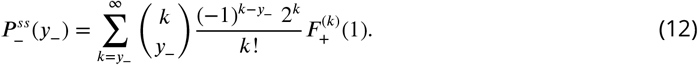

Although we have so far discussed copy numbers of the new born (*y*_+_) and most mature cells before division (*y*_−_), the calculations above may be extended to obtain the cyclo-stationary distribution *P* ^*ss*^(*y, τ*) at any intermediate cell age *τ* as shown in Sec. VI. The age-averaged distributions 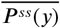 may further be obtained, given appropriate weights of cell age.

The first three cumulants associated with 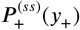 are given exactly in terms of the same coeffi-cients 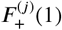 as (see SM Sec-IE ***sup (2025***)):

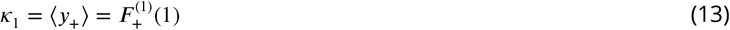

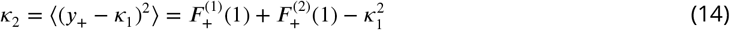

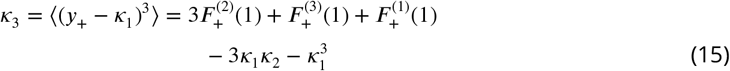

Note that below we will study the standard measures of fluctuations, namely, 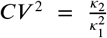 and 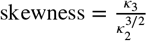 for 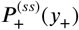 using the above Eqs. 13-15.

In summary, while for constant *t*_*s*_ = *T* the exact generating function *F*_+_(*q*) (like in Eq. 8) may be found and inverted (through derivatives) to obtain the 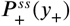, here we have shown that for more realistic random *t*_*s*_, the distribution 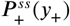 may be found directly as series sums (Eqs. 11), once the crucial coefficients 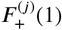 are evaluated through an exact recursion formula like Eq. 10.

## III. The Models and distributions studied

### Gene expression

We consider the basic model of constitutive gene expression (see Fig.2), in which mRNAs are produced (*m* → *m* + 1) at a rate *k*_*m*_ from the DNA template, and they degrade (*m* → *m* − 1) at a rate *γ*_*m*_. Proteins are produced from mRNAs at a rate *k*_*p*_, and they degrade (*n* → *n* − 1) at a rate *γ*_*p*_. In this work we focus on cells which have very slow protein degradation compared to the mRNA (i.e. 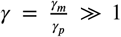). This scenario is common in yeast and bacteria, and one may treat the production of proteins effectively in bursts, ignoring the intermediate creation of mRNAs ***Yu et al. (2006); Friedman et al. (2006); Cai et al. (2006); Shahrezaei and Swain (2008***). Hence in the protein production model we study, protein copy number *n* → *n* + *r* in a burst, with the increment *r* distributed geometrically as 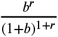. The mean burst size 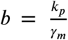 and the rate of production is indicated in Fig. 2. The degradation rate *γ*_*p*_ is taken to be finite throughout this work.

**Figure 2.**
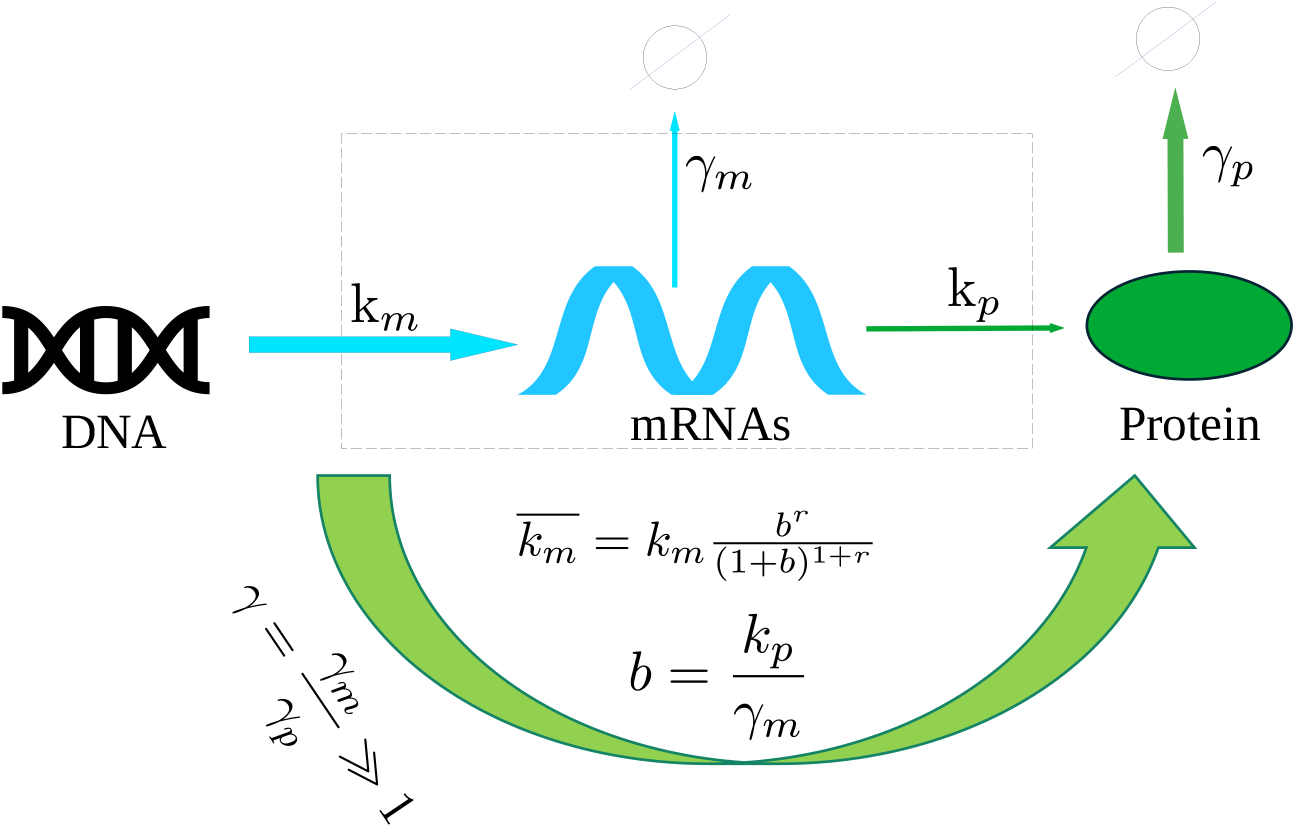
A schematic figure showing transcriptional production of mRNAs from DNA at rate *k*_*m*_ and their translation to protein at rate *k*_*p*_. They degrade at rates *γ*_*m*_ and *γ*_*p*_ respectively. In the limit of slow protein decay, i.e *γ* = *γ*_*m*_/*γ*_*p*_ *≫* 1, the protein production is bursty with an effective rate 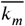 with average burst size *b*.

Note that all our mathematical results for the mRNA in this paper can be used as it is for proteins that have non-bursty production (*n* → *n* + 1) and degradation, at constant rates.

### Division time distributions

Although our results would apply to any *g*(*t*_*s*_), for concrete comparison, we study few distributions below with the same *t*_*s*_ = *T* but different *CV* ^2^. The base line case is constant ⟨*t*_*s*_⟩ = *T* with *g*(*t*_*s*_) = *δ*(*t*_*s*_ − *T*) has *CV* ^2^ = 0. The opposite extreme is the Exponential distribution 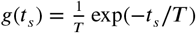 with ⟨*t*_*s*_⟩ = *T* and *CV* ^2^ = 1. The third one is the Erlang distribution

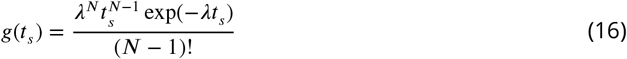

with 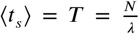 and *CV* ^2^ = 1/*N*. Note that the Erlang interpolates between the Exponential (*N* = 1) and Dirac-delta (*N* → *∞*). It is a popular distribution studied in many earlier works ***Yates et al. (2017); Antunes and Singh (2015); Perez-Carrasco et al. (2020); Beentjes et al. (2020***), but analyzed as an effective *N* step Markov process with exponentially distributed waiting times 1/*λ* ***Soltani and Singh (2016); Beentjes et al. (2020***). We would however be using it directly (like ***Antunes and Singh (2015***)) with the given form of *g*(*t*_*s*_) in Eq.16.

We would also study distributions of time *t*_*s*_, where division arises due to some threshold crossing. The Beta exponential distribution

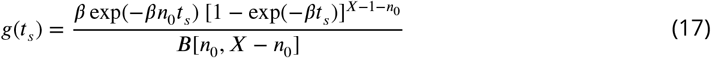

describes cell division times *t*_*s*_ when a *fixed threshold X* is crossed for the first time by either cell size or protein biomass in an autocatalytic growth process from an initial amount *n*_0_ at rate *β*. It has been applied to cell division times in bacteria C. crescentus ***Iyer-Biswas et al. (2014a***,b) ^1^. Below, we choose the initial quantity 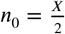 (i.e., half of the threshold), and the values of threshold *X* and growth rate *β*, to have the mean cell cycle time 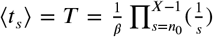 as desired in Fig. 3. The *CV* ^2^ of the Beta exponential distribution works out to be lower than the Erlang in Fig. 3, for our chosen parameters.

**Figure 3.**
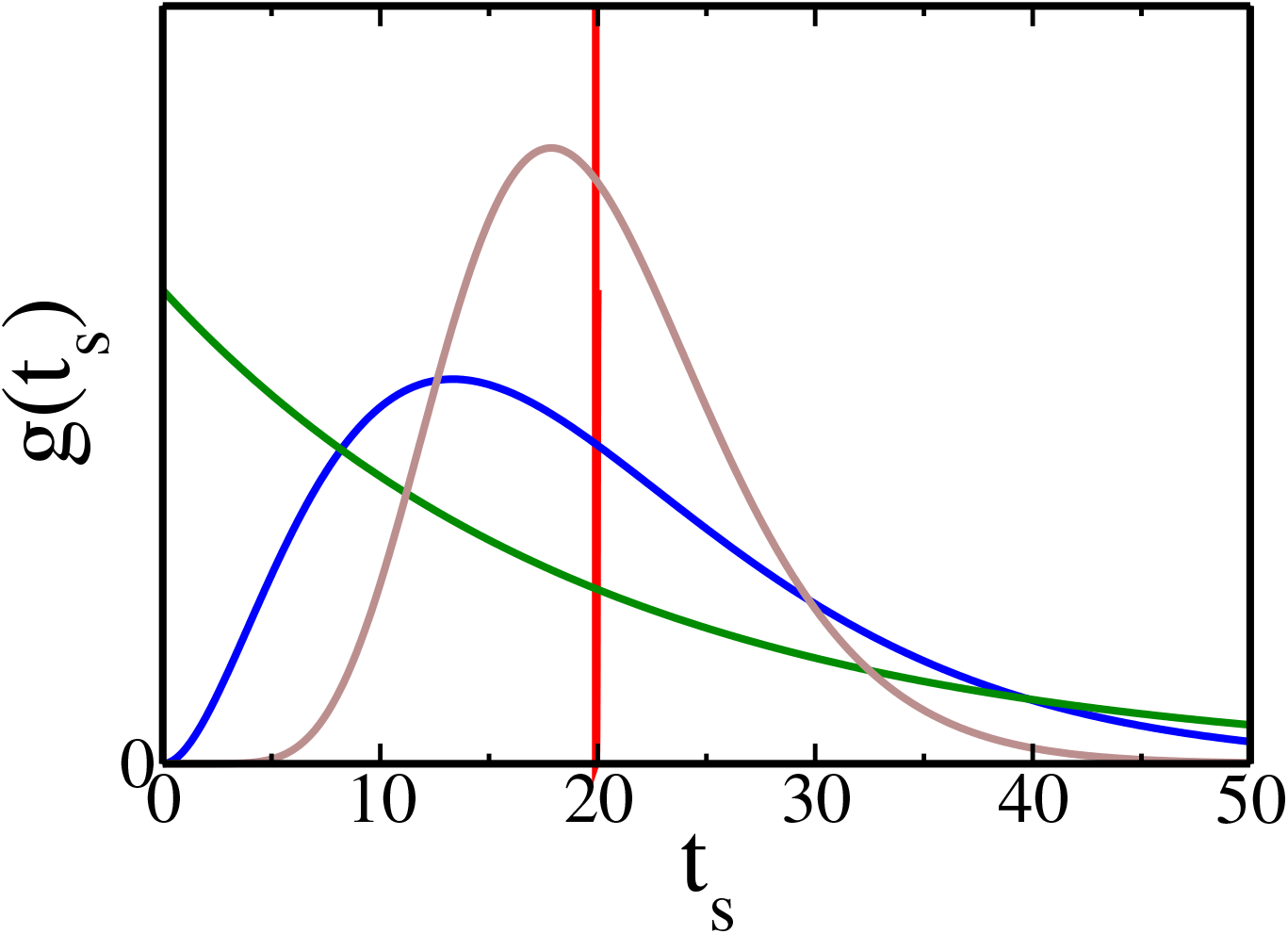
The four cell cycle time distributions *g*(*t*_*s*_) versus *t*_*s*_, used in the text: Dirac-delta (red), Beta exponential (grey) with *X* = 20 and *n*_0_ = *X*/2, Erlang (blue) with *N* = 3, Exponential (green). Other parameters are chosen so that they all have the same average ⟨ *t*_*s*_ ⟩= *T* = 20 mins. Their *CV* ^2^ values are 0, 0.104, 0.33, and, 1, respectively.

The quantity which crosses the threshold *X* to give rise to the Beta exponential distribution (Eq. 17), grows stochastically with its mean growing exponentially as *∼* exp(*βt*). Instead, if a deterministic growth of cell size is considered as *∼* exp(*βt*), but the growth rate *β* has a Gaussian variation 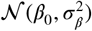 within a population of cells, then for *X* = *n*_0_/2 the effective division time distribution is

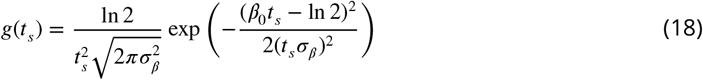

Recently, the possibility of a fluctuating (not fixed) threshold *X* has been considered ***Luo et al. (2023); Biswas and Brenner (2024***), which is crossed by an exponentially growing cell size. With a Gaussian fluctuation of ln(*X*/*n*_0_) having variance 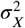, the following distribution was obtained ***Biswas and Brenner (2024***) and compared with experiments:

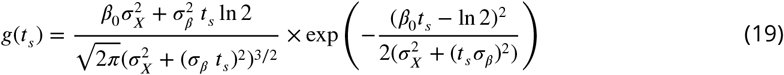

In Sec. IV we will compare the exact cyclo-stationary distributions for Eqs. 18, 19 with different *σ*_*X*_ (threshold width).

## IV. Cyclo-stationary distributions of mRNA

The Master equation for the stochastic kinetics of mRNA number *m*(*t*) within a cell cycle starting from 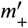 at the beginning of the cycle, is provided in SM Sec-IIA ***sup (2025***), and is solved to obtain the generating function 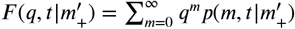 having a form same as Eq. 5:

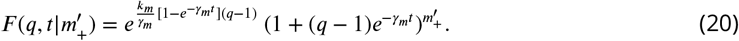

Thus here the function 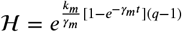. Hence the generating functions for the cyclo-stationary distributions of mRNA 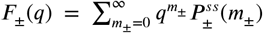 satisfy Eqs. 6 and 7 with *y*_±_ = *m*_±_. We show in SM Sec-IIB ***sup (2025***) that the series expansion of *F*_+_(*q*) about *q* = 1 (similar as Eq. 9) leads to the following recursion relation among the coefficients 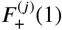 (of the form as Eq. 10):

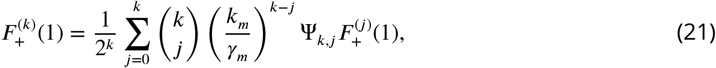

where 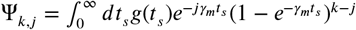.

The recursive Eq. 21 can be exactly solved for 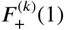 (see Eq. 39 in SM Sec-IIB ***sup (2025***)). Hence through Eq. 11 and 12, cyclo-stationary distributions 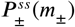 are formally solved. The expression for 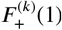 is a bit cumbersome, involving sum over subsets of integers. For the following two *g*(*t*_*s*_), simpler closed forms are obtained:

1. For *g*(*t*_*s*_) = *δ*(*t*_*s*_ − *T*), it may be shown (see details in SM Sec IIC ***sup (2025***)) that 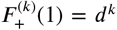 with 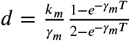. The corresponding cyclo-stationary distributions (from Eqs. 11,12) are *Poisson*:

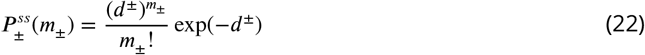

with *d*^+^ = *d* and *d*^−^ = 2*d*.
2. For Exponential distribution 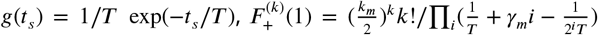 (see SM Sec-IID ***sup (2025***)) and Eqs. 11,12 lead to

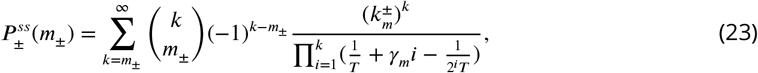

with 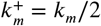 and 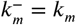. As noted above, all the analytical expressions for mRNA in this section also apply to non-bursty protein kinetics.

For other distributions in general (including Erlang and Beta Exponential), simpler formulas than Eq. 39 in SM ***sup (2025***) for 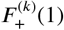 are not apparent. Alternatively, it is more efficient to obtain these coefficients numerically from the exact recursion formula Eq. 21, and use those to obtain 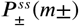 by summing the series in Eqs. 11 and 12. Such numerical summation protocol needs some care as the coefficients 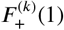 grow exponentially large with *k* and the series for 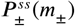 typically converge very slowly. It may be done in many ways. We suggest storing the logarithms of large numbers, using very high precision in Mathematica and, if necessary, using the Borel summation method for quicker convergence – these are discussed in Sec-IV of SM ***sup (2025***).

In Fig. 4(a), the analytically obtained 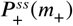 (in solid lines) are shown for the four distributions from Fig. 3. They are validated by independent data from Gillespie simulations ***Gillespie (1977***) of these models (see SM Sec.IV ***sup (2025***)). The differences of the curves in Fig. 4(a), reflect the differences in the ‘extrinsic’ factor (namely the division time statistics). Note that not just few moments but the full distributions *g*(*t*_*s*_) contribute through Ψ_*k,j*_ and 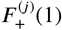 to different 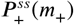.

**Figure 4.**
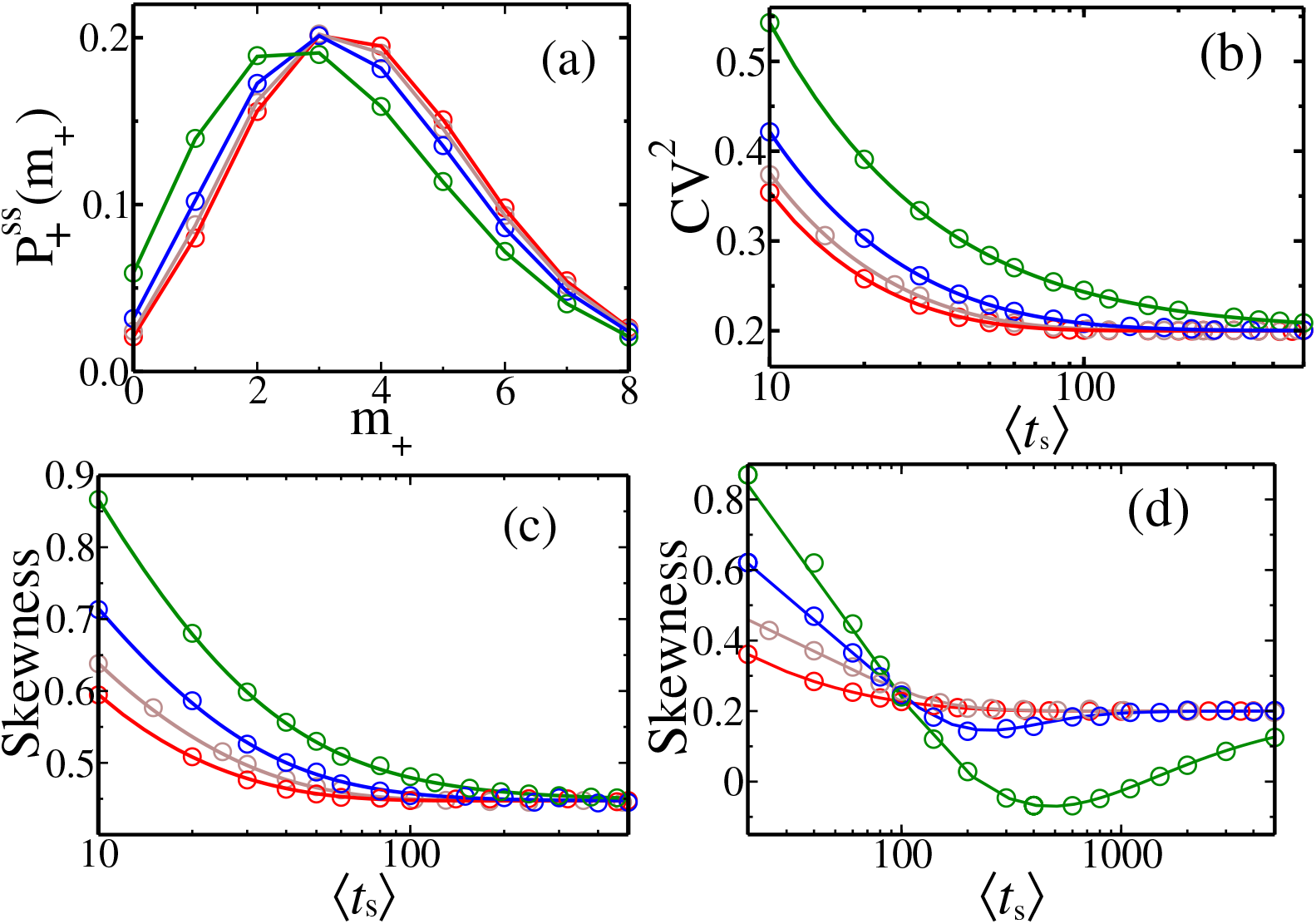
(a) Cyclo-stationary distribution 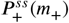 of mRNA for the four *g*(*t*_*s*_) shown in Fig.3 (corresponding colours being the same). The corresponding variation of *CV* ^2^ (b) and skewness (c) with varying ⟨*t*_*s*_⟩ are shown. Here *k*_*m*_ = 0.5, and *γ*_*m*_ = 0.05. (d) Skewness for *k*_*m*_ = 0.5, and *γ*_*m*_ = 0.01. The solid lines follow analytical formulas, while filled symbols represent KMC simulation data (using 10 × 10^7^ histories).

Further insight on cyclo-stationary fluctuations come from study of second and third order cumulants. Since the necessary quantities 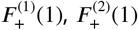, and 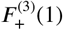 appearing in Eqs 13-15 may be exactly solved in terms of Ψ_*k,j*_ (see Eqs. 36, 37 and 38 in Sec-IIB of SM ***sup (2025***)), one may study the *CV* ^2^ and Skewness for any *g*(*t*_*s*_). The explicit formula for

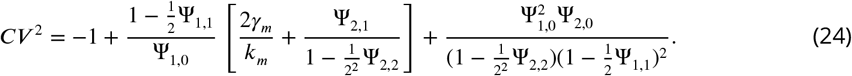

In Fig. 4(b) we see that the *CV* ^2^ of 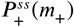 vary monotonically with mean division times ⟨*t*_*s*_⟩, and for all distributions approach the asymptotic value of 2*γ*_*m*_/*k*_*m*_ (= 0.2 in Fig. 4(b)) for large ⟨*t*_*s*_ ⟩. This follows immediately for Dirac-delta distribution from Eq. 24, as 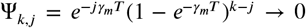 for any *j* ≠ 0, and → 1 for *j* = 0, at large *T*. For other *g*(*t*_*s*_), the times *t*_*s*_ around the mean dominate the integral of Ψ_*k,j*_ at large ⟨*t*_*s*_⟩, implying similar asymptotic values. In Fig.4(b), we observe that slower is the decrease of *CV* ^2^ of mRNA count, when higher is the *CV* ^2^ of *g*(*t*_*s*_) (see the hierarchy in Fig 3), as slower are the corresponding approaches of Ψ_*k,j*_ to the asymptotic values. The *CV* ^2^ can never be non-monotonic with ⟨*t*_*s*_⟩, since the contributing terms 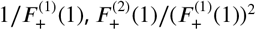 add with the same sign and each monotonically decrease.

In Fig.4(c), we see a similar asymptotic approach of Skewness to a value 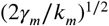(= 0.447 in the figure), for any *g*(*t*_*s*_). This follows from the formula of Skewness in Eq. 57 in SM ***sup (2025***), based on the arguments given above that Ψ_*k,j*_ → 0 for *j* ≠ 0, and → 1 for *j* = 0 at large ⟨*t*_*s*_⟩ for any *g*(*t*_*s*_). A striking feature of the Skewness formula is that it may exhibit non-monotonic behavior if the degradation rate (i.e., *γ*_*m*_) is sufficiently small. Observe although curves in Fig.4(c) are monotonic for *γ*_*m*_ = 0.05, they are non-monotonic in Fig.4(d) for smaller *γ*_*m*_ = 0.01 before asymptotically flattening. The reason is that, in contrast to *CV* ^2^, in the Skewness formula, some of the terms dependent on *g*(*t*_*s*_) have a plus sign while some have a minus sign. Hence if the group of terms with the minus sign are relatively slower in attaining their asymptotic value in comparison to the group of terms with a positive sign, the value of Skewness can get depressed and then again rise as a function of ⟨*t*_*s*_⟩ (as in Fig.4(d)) – this effect will be magnified if *γ*_*m*_ is small and *CV* ^2^ of *g*(*t*_*s*_) is high, both delaying the asymptotics. We will show below this interesting feature in Skewness is also present for the Skewness of protein counts, due to similar reasons.

We have also studied the analytical probability distributions 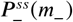 of mRNA count just before cell division and compared with simulation data – see Fig. 1 in SM ***sup (2025***), for four different *g*(*t*_*s*_).

## V. Cyclo-stationary distributions of protein

In Sec-III of supplementary text ***sup (2025***), the Master equation for the bursty production and degradation kinetics (see Fig. 2) of protein number *n*(*t*), starting from 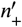 at the beginning of the cycle, is shown. The corresponding probability distribution 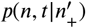 has a generating function 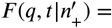

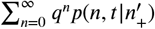 ***Shahrezaei and Swain (2008***):

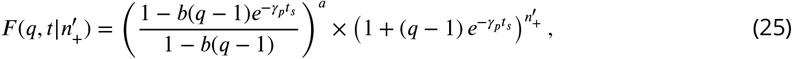

where *a* = *k*_*m*_/*γ*_*p*_ (see SM Sec IIIA ***sup (2025***)). The above equation has the same form as Eq. 5 with the function 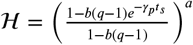. Hence Eqs. 6 and 7 are satisfied by the generating functions 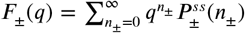, for the cyclo-stationary distributions 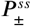 of proteins. We show in SM SecIIIB ***sup (2025***) that the series expansion of *F*_+_(*q*) about *q* = 1 (see Eq. 9) yields the following exact recursion relation for 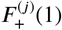:

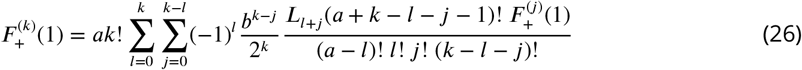

Where 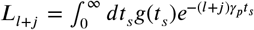, a function of (*l* + *j*)*γ*_p_, is the Laplace transform of the distribution *g*(*t*_*s*_). The Eq. 26 is the key exact result – it is reducible to the form in Eq. 10 by combining terms. The coefficients 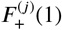 can be enumerated using Eq. 26 once the Laplace transform of the random cell cycle distribution *g*(*t*_*s*_) is known, and those in turn lead to the analytical cyclo-stationary distributions 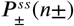 through Eqs. 11 and 12 (see the protocol in Sec-IV of SM ***sup (2025***)). In Fig 5(a) and (d) the analytical protein distributions 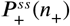 and 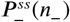 (in solid lines) are well matched by simulation data (in symbols).

**Figure 5.**
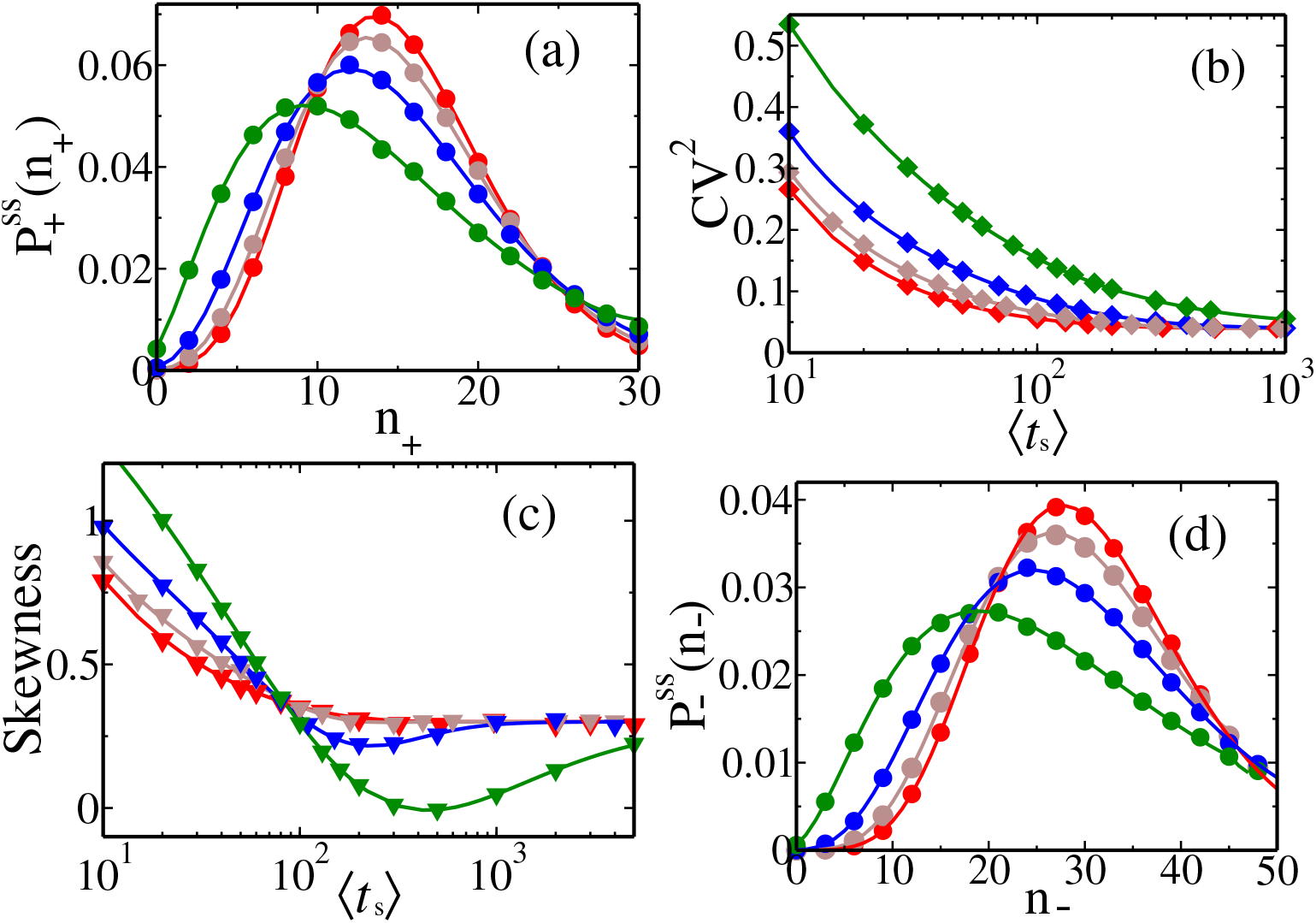
(a) Cyclo-stationary distribution 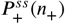 of protein count, for the four *g*(*t*_*s*_) shown in Fig.3 (corresponding colours being the same). The corresponding variation of *CV* ^2^ (b) and Skewness (c) with varying ⟨*t*_*s*_⟩ are shown. (d) The distribution 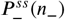 just before division. Here *k* = 0.5, *b* = 2, and *γ* = 0.01. The solid lines follow analytical formulas, while empty symbols represent KMC simulation data (using 5 × 10 histories).

The exact 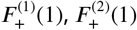, and 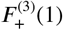, are provided in Eqs. 63, 64 and 65 in SM Sec-IIIC ***sup (2025***), in terms of *L*_*k*_. The exact *CV* ^2^ and Skewness for any *g*(*t*_*s*_) (using Eqs 13-15) for the protein count at cell birth, follow from those. For convenience of user, we provide below:

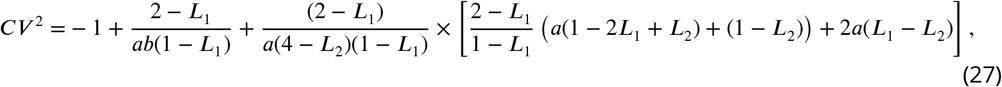

Like in the case of mRNA, the *CV* ^2^ of *n*_+_ are hierarchical, with respect to the degree of fluctuations in *g*(*t*_*s*_) (Fig 5(b)), but they all approach a common asymptotic value. As all *L*_*j*_ → 0 at large ⟨*t*_*s*_⟩ for any *g*(*t*_*s*_), from Eq. 27, we conclude that *CV* ^2^ → (*b* + 2)/*ab* (= 0.04 in the figure). Here too the additive terms 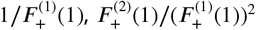 in *CV* ^2^ each monotonically decrease, implying *CV* ^2^ to be always monotonic with ⟨*t*_*s*_⟩. In Sec. VI, we analyze the contributions of intrinsic and extrinsic factors separately to *CV* ^2^ in Eq. 27.

Since the degradation rate *γ*_*p*_ of proteins are typically low, the terms in the Skewness expression will take long time to reach their asymptotic value 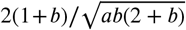 (= 0.3 in Fig. 5(c)) which follows by setting *L*_*j*_ → 0 in Eq. 95 of SI ***sup (2025***). Because of the relative difference of times of approach to the asymptotics of the groups of positive and negative terms (Eq. 95 of SI ***sup (2025***)), we have a non-monotonic behavior in the Skewness as a function of ⟨*t*_*s*_⟩ – see Fig. 5(c).

## VI. Various extensions of the above results

### A. Cyclo-stationary distributions at any cell age *τ*

Although so far we discussed the cyclo-stationary distribution of copy numbers in the new-born cells 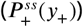 and cells prior to division 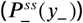, the cyclo-stationary distribution *P* ^*ss*^(*y, τ*) of cells at any arbitrary ‘age’ *τ* before the next cell division, may be derived by using the 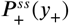 as follows:

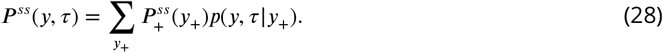

Using its generating function 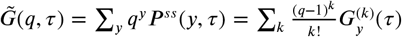, we derive (see SM Sec V ***sup (2025***))

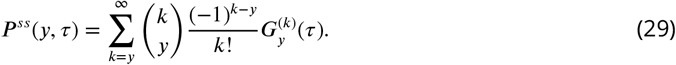

For the mRNAs (i.e *y* ≡ *m*), with *F* ^(*k*)^(1) from Eq. 21,

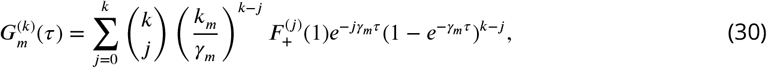

while for proteins (i.e *y* ≡ *n*) with *F* ^(*k*)^(1) from Eq. 26,

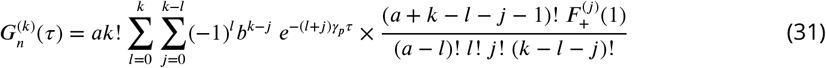

See SM ***sup (2025***)) for details of the calculation. Thus we have the analytical formulas for *P* ^*ss*^(*y, τ*) given by Eq 29, along with Eqs 30, 31. Note that when 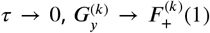 and hence 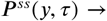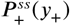 (Eq. 11) as expected. In Fig. 6, we show the plots of the theoretical *P* ^*ss*^(*m, τ*) and *P* ^*ss*^(*n, τ*) at three different ages, using the Erlang distribution for the *g*(*t*_*s*_), and they match very well with the simulation data.

**Figure 6.**
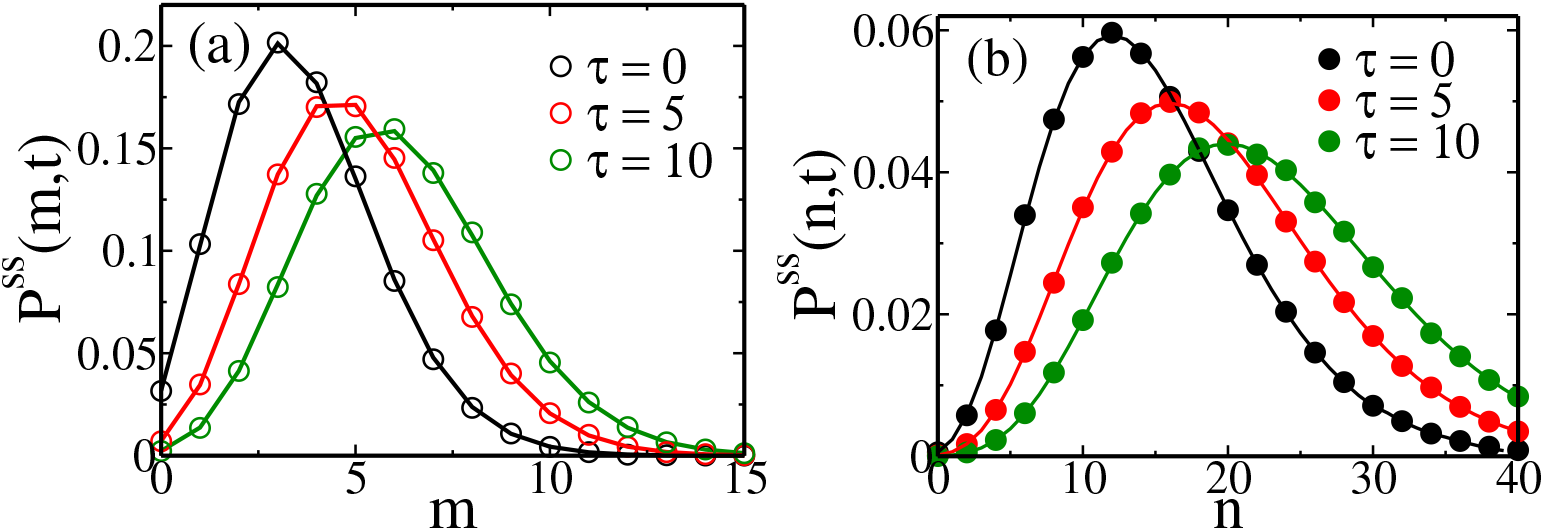
Cyclo-stationary distributions *P* ^*ss*^(*m, τ*) of mRNA (m) and *P* ^*ss*^(*n, τ*) of protein (n), at three different cell ages (*τ* = 0, 5 and 10 mins). The solid lines are analytical theory, while symbols are simulation data. The *g*(*t*_*s*_) follows the Erlang distribution from Fig. 3. The parameters for mRNA are *k*_*m*_ = 0.5, *γ*_*m*_ = 0.05, and protein are *k*_*m*_ = 0.5, *b* = 2, and *γ*_*p*_ = 0.01.

### B. Age-averaged cyclo-stationary distributions

Distributions averaged over cell age, would require the relative weight of the cell-age *ϕ*(*τ*). In some studies, uniform weight of all ages was assumed in ***Soltani and Singh (2016); Soltani et al. (2016); Beentjes et al. (2020***), corresponding to a single lineage.

In a large population of exponentially growing cells, but with ‘constant’ cell division times, the age equation 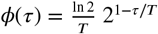, and has been used in various works ***Berg (1978); Rigney (1979); Beentjes et al. (2020); Wang et al. (2023***).

However, we need the age distribution *ϕ*(*τ*) for random cell division times with a given *g*(*t*_*s*_). The formula for that was derived by Powell ***Powell (1956***): 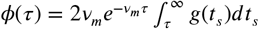 with *ν* given implicitly by 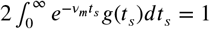.Thus one needs to first calculate the appropriate *ν*_*m*_ and *ϕ*(τ) given a *g* (*t*_*s*_). For example for Erlang distribution (Eq. 16), 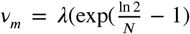 and 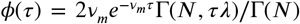. Then the age-averaged distribution is 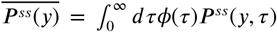, which may be evaluated using *P* ^*ss*^(*y, τ*) (Eq. 29). For the Erlang distribution, the curves are shown in Fig. 7 for mRNA and protein.

**Figure 7.**
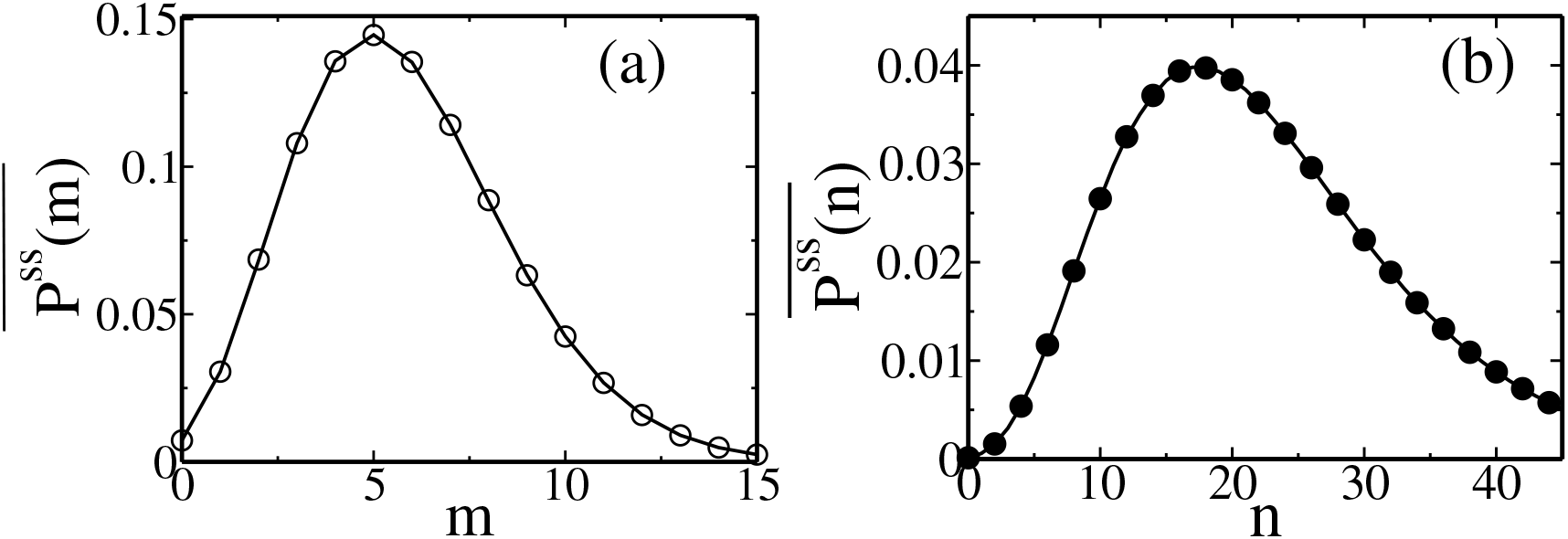
Age-averaged cyclo-stationary distribution (a) 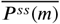 (of mRNA) and (b) 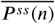 (of protein) for age distribution *ϕ*(*τ*) (see text) corresponding to the Erlang form (with *N* = 3) of *g*(*t*_*s*_). The solid lines are analytical theory, while symbols are simulation data. The *g*(*t*_*s*_) follows the Erlang distribution in Fig. 3. The parameters for mRNA are *k*_*m*_ = 0.5, *γ*_*m*_ = 0.05, and protein are *k*_*m*_ = 0.5, *b* = 2, and *γ*_*p*_ = 0.01.

### C. Comparing the contributions to noise in protein copy number, from intrinsic and extrinsic sources

In the expression of *CV* ^2^ for proteins at cell birth (Eq.27) the noise from gene expression, binomial partitioning, as well as random cell division times, all contribute. Note that mean ⟨*n*_+_⟩ = *ab*(1 − *L*_1_)/(2 − *L*_1_) is independent of whether the processes are deterministic or stochastic. If we wish to exclude the contribution of the partitioning noise, then the Binomial distribution may be replaced by a delta function (in Eq. 3) such that *n*_+_ = *n*_−_/2 in every cycle. A separate calculation done for this case in SM Sec VI ***sup (2025***), shows that (*q* − 1)/2 in Eq.6 for the generating function is replaced by 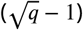, and subsequently yields a

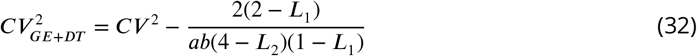

where *CV* ^2^ is the total noise (from Eq. 27). The 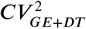 has a contribution from gene expression (GE) and division time (DT) randomness. Thus the noise from Binomial partitioning (BP) is

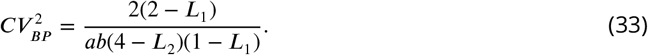

Note 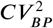 is dependent on *g*(*t*_*s*_), although mildly – see Table 1, for the different cases. If in addition to removal of partitioning noise (i.e. setting *n*_+_ = *n*_−_/2), further the gene expression is also deterministic, then protein number at any time 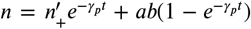 (with initial count 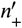). In this case, the argument of *F* (.) on the right hand side of Eq. 6 is replaced by 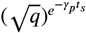 (see SM Sec VI ***sup (2025***)), and the calculation gives

**Table 1.** Noise in *n*_+_, for ⟨*t*_*s*_⟩ = 20 mins, *a* = 50, *b* = 2

| $g(t_s)$ | $CV^2$ | $CV_{BP}^2$ | $CV_{DT}^2$ | $CV_{GE}^2$ |
| --- | --- | --- | --- | --- |
| Dirac-Delta | 0.149 | 0.039 | 0 | 0.110 |
| Beta-Exponential | 0.174 | 0.039 | 0.025 | 0.110 |
| Erlang (N=3) | 0.229 | 0.040 | 0.078 | 0.111 |
| Exponential | 0.372 | 0.042 | 0.217 | 0.113 |

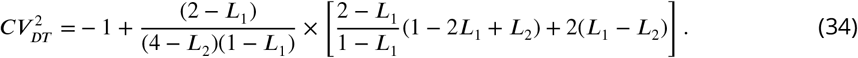

The 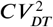 is purely the extrinsic noise contribution coming from random variations in cell division times – hence if *g* = *δ*(*t*_*s*_ − *T*), then 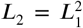, and hence 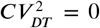 in Eq.34 as expected. Also see Table 1 for other cases. The *CV* ^2^ in Eq. 27 is in excess of 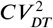 (Eq.34) by two terms. Comparing with Eq. 32, we have noise contribution from gene expression to total *CV* ^2^ as

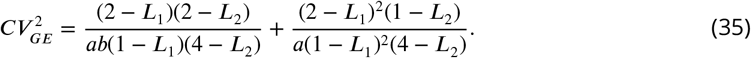

Note 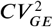 is dependent on *g*(*t*_*s*_), although mildly – see Table 1, for the different cases.

Finally percentage contributions of 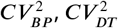, and 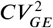 to the total *CV* ^2^ depends on the *g*(*t*_*s*_). From Table 1, for the Beta exponential, they are respectively, 23%, 14%, and 63%, for the Erlang, they are respectively, 18%, 34%, and 48%, and for the Exponential, they are respectively, 12%, 58%, and 30%. Thus as division time noise rises, the relatively higher contribution of gene expression noise is offset by it.

Here we have presented the results for protein, but following the same type of calculations, one may derive corresponding results for the mRNA and compare with the full *CV* ^2^ from Eq.24.

### D. Division time distributions due to fluctuating threshold, and effect on cyclostationary protein statistics

In Sec. III, we mentioned division time distributions arising from threshold crossing scenarios – Eq.18 due to growth rate heterogeneity but fixed threshold, and Eq. 19 due to both growth rate heterogeneity and threshold fluctuations. Partly to demonstrate the applicability of our results to these cases too, and partly to see the effect of threshold fluctuations, we study them here.

In Fig.8(a), for chosen *β*_0_ = 0.035 min^−1^, *σ*_*β*_ = 0.0052 min^−1^ (with mean division time ≈ 20 mins), three *g*(*t*_*s*_) distributions are shown with different widths *σ*_*X*_ of the size threshold in Eq. 19 – parameters are in the ballpark of ref ***Biswas and Brenner (2024***). The corresponding cyclo-stationary protein distributions 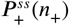 in new born cells for the three cases are shown in Fig.8(b). The copy number distributions broadens relatively slowly as threshold fluctuations (*σ*_*X*_) rise and *g*(*t*_*s*_) quickly becomes broader.

**Figure 8.**
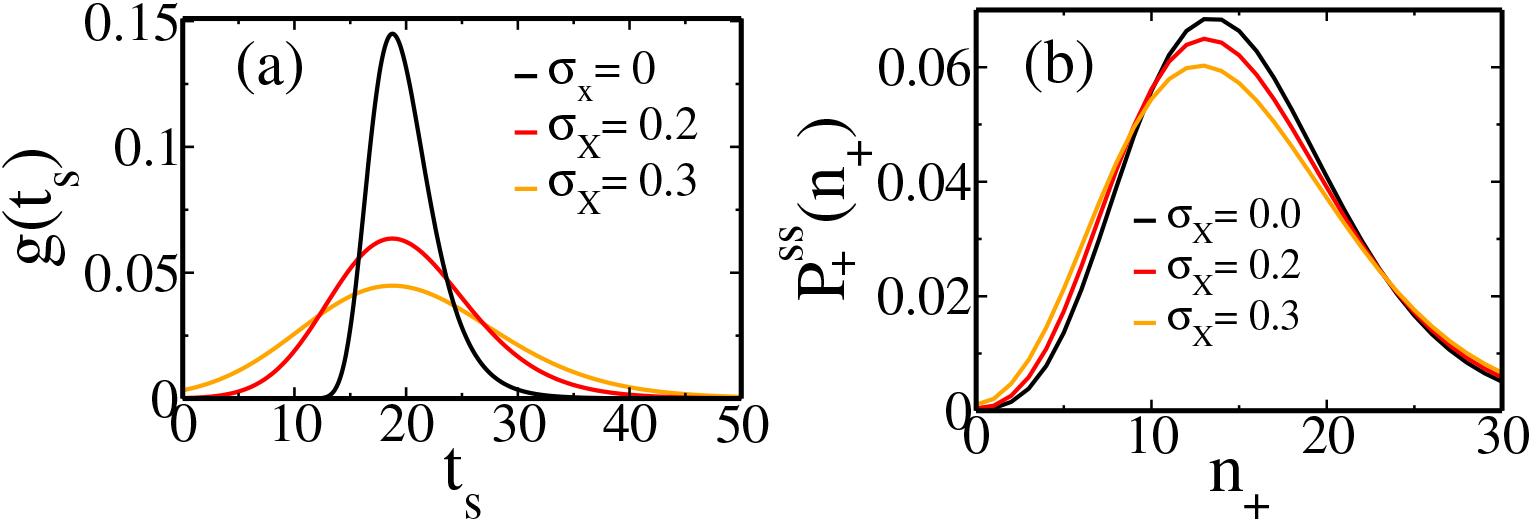
(a) Plots of different *g*(*t*_*s*_) following Eq.19, with various values of *σ*_*X*_ as shown in the figure. We have *β*_0_ = 0.035 min^−1^, *σ*_*β*_ = 0.0052 min^−1^ for all of them. (b) The corresponding cyclo-stationary distributions 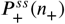 (following the same colors as in (a)) are shown. The parameters for protein gene expression are *b* = 2, *k*_*m*_ = 0.3, and *γ*_*p*_ = 0.01.

### E. Effect of correlations in cell cycle times

For deriving the basic recursive equations involving the cyclo-stationary distributions (Eq. 3 and 4) we had assumed that the random cell cycle times are uncorrelated, i.e. 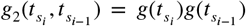. Given the complexities of cell-cycle control, this assumption may have to be relaxed. For example, recent work on specific human cell lines shows random cell-cycle times that are not correlated between the mother and daughter cells ***Chakrabarti et al. (2018***). Interestingly, data showed modest correlations between the cell-cycle times of daughter cells, but this correlation is lost between a cousin pair of cells ***Chakrabarti et al. (2018***). Here we study, through Kinetic Monte Carlo simulations, the effect of correlated cell cycle times. Correlation is achieved by choosing the time of *i*^*th*^ (*i* > 1) cycle 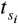 in terms of the (*i* − 1)^*th*^ cycle time 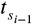 in the following way:

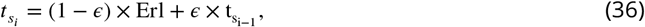

where Erl is a random time chosen for the *i*^*th*^ cycle, following the Erlang distribution in Fig. 3. For the first cycle 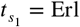. By construction, the mean of the correlated 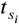 distribution remains the same as the Erlang distribution. Note that by tuning the parameter *ϵ* from zero to positive values, we can increase the degree of correlation. In Fig. 9(a) and (b), we show the corresponding cyclo-stationary distributions of mRNA and protein. With a rise of *ϵ*, both the distributions (in symbols) show some deviation (although not significant) from the uncorrelated analytical distributions (solid lines) that we have derived in this paper. The departure is more pronounced for protein than mRNA.

**Figure 9.**
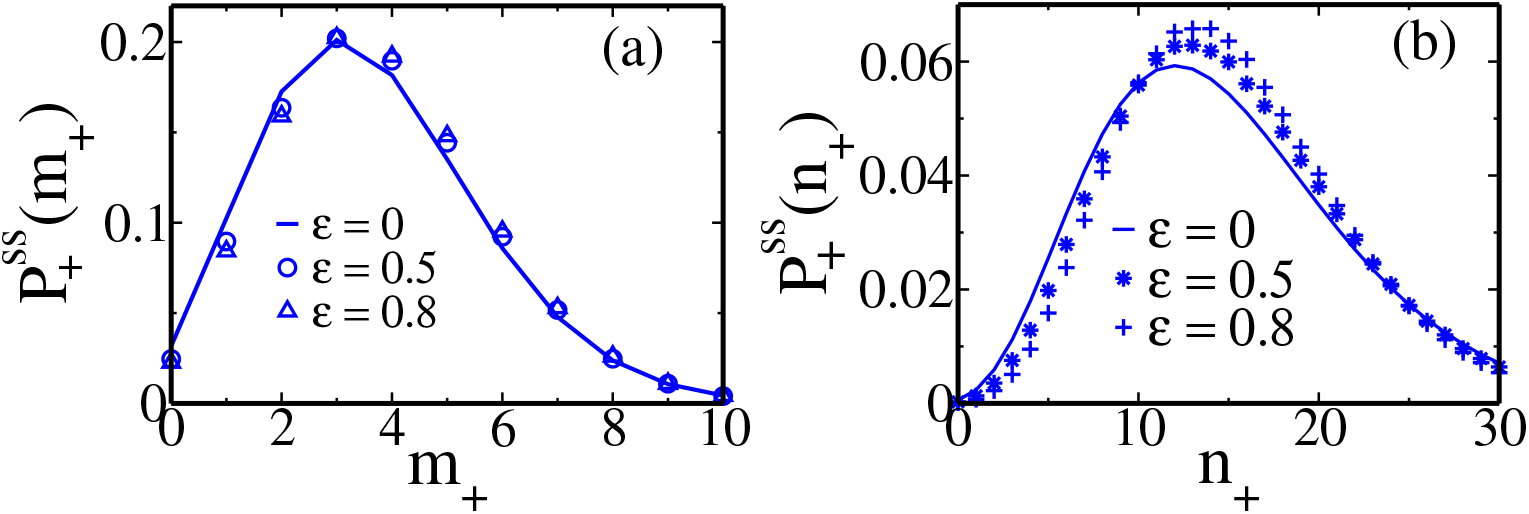
Effect of correlation in cell cycle times on the cyclo-stationary distributions of (a) mRNA and (b) protein. The analytical predictions in the uncorrelated case are in solid lines. The symbols are from KMC simulations with various degrees of correlation indicated by the parameter *ϵ*.

## VII. Concluding discussions

Analysis of single-cell transcriptomic and proteomic data requires an understanding of stochastic cellular processes that influence the variability of copy numbers of mRNA and proteins from cell to cell. Years of theoretical work on various models of gene expression, along with additional complexities of partitioning noise during division, dynamic cell growth, and gene duplication, have already enriched the means for analyzing the data. The key contribution of this paper is to present a method to incorporate the non-negligible aspect of noise in cell cycle times, within the evolving broader picture.

We have studied theoretically copy number statistics in cells obtained after many cycles of cell division, each involving binomial partitioning of copy numbers, when a cyclo-stationary condition has been attained. For any random (but uncorrelated) cell division times, we have presented a method to obtain exact series representations of the distributions of copy numbers. This is a significant theoretical advancement, as analytical solutions were known only for deterministic division times and a specific type of random time distribution (namely the Erlang). Moreover, treatments of the Erlang distribution relied on the steady state assumption of every cell cycle stage, and hence could only get cell age-averaged statistics. Our method here makes no such assumption and gives exact distributions at any cell age, and also leaves the scope of obtaining the age-averaged result using the suitable cell age distribution ***Powell (1956***). We have demonstrated that mild correlations in division times of successive cycles would not cause a strong departure from the analytical results obtained for the uncorrelated case. Extension of the current work by developing efficient summation methods of the formal series we present, would be a fruitful direction to pursue in future.

The following results followed by the consequences random cell cycle times. Along with the general series forms of cyclo-stationary distributions of mRNA and proteins for any random cell cycle distribution, we specifically showed that for fixed cycle times, the mRNA distribution is Poisson. Thus, the Poisson form, well-known for constitutive transcription, stays preserved after accounting for partitioning noise as long as the cell division time is a constant. Hence, the departure of the mRNA distribution from Poisson is a signature of the effect of cell cycle time variability (in cases where promoter regulation is absent). Next, the Skewness has non-monotonic variation with mean division time, if fluctuations in division time are strong, and the degradation rate of copy numbers is weak. This behavior of Skewness may serve as a signature of high variability of cell cycle times. In comparison, we found that *CV* ^2^ never exhibits non-monotonicity, and it is high in consonance with higher fluctuations in cell division times. Splitting the net *CV* ^2^ into contributions from intrinsic and extrinsic sources, we showed (in Table 1) that with rising fluctuations in division times, its contribution may be substantial (rising to around 50%) in comparison to noise due to gene expression and binomial partitioning.

Although the models of gene expression that we analyze are the basic ones for constitutive production, they demonstrate the new method with clarity. Further complexity in models of gene expression have to be introduced through the corresponding generating functions (analogous to Eq. 5). An important way to extend the analysis would be to directly consider the impact of dynamic changes in gene dosage and size on expression levels as the cell transitions through different cellcycle stages ***Padovan-Merhar et al. (2015***). This can be done in either models with population counts with binomial partitioning at division ***Beentjes et al. (2020); Cao and Grima (2020***), or alternatively model concentration of gene products, where the average concentration remains the same after division ***Friedman et al. (2006); Jia et al. (2022); Zhang et al. (2024); Rijal et al. (2022***). Each of these extensions may follow the core idea presented here, but is expected to be a lengthy and interesting calculation, and hence left for future study.

We made an interesting connection between two entirely different biophysical problems – namely, the problem of cell division and that of synaptic vesicle release triggered by action potentials ***Rijal et al. (2024***). The underlying mathematical structure of these two problems is similar, as both involve repeated cycles of growth (or replenishment) and partitioning (or release). Yet the specific results are different, as the details of the process of gene expression differ from those of the stochastic docking of synaptic vesicles.

Stochastic gene expression plays a critical role in a number of biologically and biomedically significant processes. Notable examples include stochastic cell fate determination in both bacteria and multicellular organisms ***Losick and Desplan (2008); Carniol et al. (2004***), spontaneous prophage induction in bacteria ***Miyazaki et al. (2012); Little and Michalowski (2010***), and the random expression of proteins that confer antibiotic resistance in bacteria ***Balaban et al. (2004***) or chemotherapy resistance in cancer cells ***Schuh et al. (2020); Shaffer et al. (2020***). Analytical models of protein expression are thus potentially very useful for analyzing noise-driven processes in biology. Previous models for the distribution of mRNA and protein levels in cells did not fully account for the contributions from randomness in cell division times in conjunction with binomial partitioning. In this paper, we incorporate these sources of extrinsic noise into an analytical exact theory of mRNA and protein gene expression. We anticipate that the results would have wide applicability in biomedical and biological research for systems where noise in cell cycle duration plays a functional role.

## Supporting information

Manuscript

## Acknowledgments

DD acknowledges the visitor program of MPI-PKS Dresden (where a part of this work was done in summer 2024), and thanks K. Rijal and Madan Rao for discussions. SYA acknowledges IIT Bombay for financial support through the institute post-doctoral fellowship. AS acknowledges support from NIH-NIGMS via grant R35GM148351.

## Footnotes

1 Interestingly, the first passage times of threshold crossing of bacterial population size undergoing autocatalytic growth follows the Beta Exponential distribution, as was shown by Dellbruck years before ***Delbrück (1940***).

## Notes

### Competing Interest Statement

The authors have declared no competing interest.

## References

Supplementary Material; 2025. url will be inserted by publisher.

Antunes D, Singh A. Quantifying gene expression variability arising from randomness in cell division times. J Math Biol. 2015; 71:437–463. doi: 10.1007/s00285-014-0811-x.

Balaban NQ, Merrin J, Chait R, Kowalik L, Leibler S. Bacterial Persistence as a Phenotypic Switch. Science. 2004; 305(5690):1622–1625. https://www.science.org/doi/abs/10.1126/science.1099390, doi: 10.1126/sci-ence.1099390.

Beentjes CHL, Perez-Carrasco R, Grima R. Exact solution of stochastic gene expression models with bursting, cell cycle and replication dynamics. Phys Rev E. 2020 Mar; 101:032403. https://link.aps.org/doi/10.1103/PhysRevE.101.032403, doi: 10.1103/PhysRevE.101.032403.

Berg OG. A model for the statistical fluctuations of protein numbers in a microbial population. J Theor Biol. 1978; 71(4):587–603. https://www.sciencedirect.com/science/article/pii/0022519378903260, doi: 10.1016/0022-5193(78)90326-0.

Bertaux F, Marguerat S, Shahrezaei V. Division rate, cell size and proteome allocation: impact on gene expression noise and implications for the dynamics of genetic circuits. R Soc Open Sci. 2018; 5(3):172234. https://royalsocietypublishing.org/doi/abs/10.1098/rsos.172234, doi: 10.1098/rsos.172234.

Bird AD, Wall MJ, Richardson MJ. Bayesian Inference of Synaptic Quantal Parameters from Correlated Vesicle Release. Front Comput Neurosci. 2016; 10:116. doi: 10.3389/fncom.2016.00116.

Biswas K, Brenner N. Universality of phenotypic distributions in bacteria. Phys Rev Res. 2024 May; 6:L022043. https://link.aps.org/doi/10.1103/PhysRevResearch.6.L022043, doi: 10.1103/PhysRevResearch.6.L022043.

Bokes P, King JR, Wood AT, Loose M. Exact and approximate distributions of protein and mRNA levels in the low-copy regime of gene expression. J Math Biol. 2012; 64:829–854. doi: 10.1007/s00285-011-0433-5.

Cadart C, Monnier S, Grilli J, Sáez PJ, Srivastava N, Attia R, Terriac E, Baum B, Cosentino-Lagomarsino M, Piel M. Size control in mammalian cells involves modulation of both growth rate and cell cycle duration. Nat Commun. 2018; 9(1):3275. doi: 10.1038/s41467-018-05393-0.

Cai L, Friedman N, Xie XS. Stochastic protein expression in individual cells at the single molecule level. Nature. 2006 Mar; 440(7082):358–362. doi: 10.1038/nature04599.

Campos M, Surovtsev IV, Kato S, Paintdakhi A, Beltran B, Ebmeier SE, Jacobs-Wagner C. A Constant Size Extension Drives Bacterial Cell Size Homeostasis. Cell. 2014; 159(6):1433–1446. https://www.sciencedirect.com/science/article/pii/S0092867414014998, doi: 10.1016/j.cell.2014.11.022.

Cao J, Packer JS, Ramani V, et al. Comprehensive single-cell transcriptional profiling of a multicellular organism. Science. 2017; 357(6352):661–667. https://www.science.org/doi/abs/10.1126/science.aam8940, doi: 10.1126/science.aam8940.

Cao Z, Grima R. Analytical distributions for detailed models of stochastic gene expression in eukaryotic cells. Proc Natl Acad Sci USA. 2020; 117(9):4682–4692. https://www.pnas.org/doi/abs/10.1073/pnas.1910888117, doi: 10.1073/pnas.1910888117.

Carniol K, Eichenberger P, Losick R. A Threshold Mechanism Governing Activation of the Developmental Regulatory protein ???? in Bacillus subtilis. J Biol Chem. 2004; 279(15):14860–14870. doi: 10.1074/jbc.M314274200.

Chakrabarti S, Paek AL, Reyes J, Lasick KA, Lahav G, Michor F. Hidden heterogeneity and circadian-controlled cell fate inferred from single cell lineages. Nat Commun. 2018; 9(1):5372. doi: 10.1038/s41467-018-07788-5.

Chang CA, Jen J, Jiang S, et al. Ontogeny and vulnerabilities of Drug-Tolerant Persisters in HER2+ Breast Cancer. Cancer Discov. 2022; 12(4):1022–1045. doi: 10.1158/2159-8290.CD-20-1265.

Cookson NA, Cookson SW, Tsimring LS, Hasty J. Cell cycle-dependent variations in protein concentration. Nucleic Acids Res. 2009 12; 38(8):2676–2681. https://doi.org/10.1093/nar/gkp1069, doi: 10.1093/nar/gkp1069.

Delbrück M. Statistical Fluctuations in Autocatalytic Reactions. J Chem Phys. 1940 01; 8(1):120–124. https://doi.org/10.1063/1.1750549, doi: 10.1063/1.1750549.

Deloupy A, Sauveplane V, Robert J, Aymerich S, Jules M, Robert L. Extrinsic noise prevents the independent tuning of gene expression noise and protein mean abundance in bacteria. Sci Adv. 2020; 6(41):eabc3478. https://www.science.org/doi/abs/10.1126/sciadv.abc3478, doi: 10.1126/sciadv.abc3478.

Dessalles R, Fromion V, Robert P. Models of protein production along the cell cycle: An investigation of possible sources of noise. PLoS One. 2020 01; 15(1):1–25. https://doi.org/10.1371/journal.pone.0226016, doi: 10.1371/journal.pone.0226016.

El Meouche I, Jain P, Jolly MK, Capp JP. Drug tolerance and persistence in bacteria, fungi and cancer cells: Role of non-genetic heterogeneity. Transl Oncol. 2024; 49:102069. https://www.sciencedirect.com/science/article/pii/S1936523324001967, doi: 10.1016/j.tranon.2024.102069.

El Meouche I, Siu Y, Dunlop MJ. Stochastic expression of a multiple antibiotic resistance activator confers transient resistance in single cells. Sci Rep. 2016; 6(1):19538. doi: 10.1038/srep19538.

Fantes P, Nurse P. Control of cell size at division in fission yeast by a growth-modulated size control over nuclear division. Exp Cell Res. 1977; 107(2):377–386. doi: 10.1016/0014-4827(77)90359-7.

Farquhar KS, Charlebois DA, Szenk M, Cohen J, Nevozhay D, Balázsi G. Role of network-mediated stochasticity in mammalian drug resistance. Nat Commun. 2019; 10(1):2766. doi: 10.1038/s41467-019-10330-w.

Friedman N, Cai L, Xie XS. Linking Stochastic Dynamics to Population Distribution: An Analytical Framework of Gene Expression. Phys Rev Lett. 2006 Oct; 97:168302. https://link.aps.org/doi/10.1103/PhysRevLett.97.168302, doi: 10.1103/PhysRevLett.97.168302.

Gardiner CW. Handbook of Stochastic Methods for Physics, Chemistry and the Natural Sciences, vol. 115. Springer-Verlag, Berlin, Heidelberg, New York; 1985. https://onlinelibrary.wiley.com/doi/abs/10.1002/bbpc.19850890629, doi: 10.1002/bbpc.19850890629.

Ghusinga KR, Vargas-Garcia CA, Singh A. A mechanistic stochastic framework for regulating bacterial cell division. Sci Rep. 2016; 6(1):30229. doi: 10.1038/srep30229.

Gillespie DT. Exact stochastic simulation of coupled chemical reactions. J Phys Chem. 1977 Dec; 81(25):2340– 2361. https://doi.org/10.1021/j100540a008, doi: 10.1021/j100540a008.

Golding I, Paulsson J, Zawilski SM, Cox EC. Real-Time Kinetics of Gene Activity in Individual Bacteria. Cell. 2005; 123(6):1025–1036. doi: 10.1016/j.cell.2005.09.031.

Hawkins ED, Markham JF, McGuinness LP, Hodgkin PD. A single-cell pedigree analysis of alternative stochastic lymphocyte fates. Proc Natl Acad Sci USA. 2009; 106(32):13457–13462. doi: 10.1073/pnas.0905629106.

Huh D, Paulsson J. Non-genetic heterogeneity from stochastic partitioning at cell division. Nat Genet. 2011; 43(2):95–100. doi: 10.1038/ng.729.

Huh D, Paulsson J. Random partitioning of molecules at cell division. Proc Natl Acad Sci USA. 2011; 108(36):15004–15009. https://www.pnas.org/doi/abs/10.1073/pnas.1013171108, doi: 10.1073/pnas.1013171108.

Iyer-Biswas S, Crooks GE, Scherer NF, Dinner AR. Universality in Stochastic Exponential Growth. Phys Rev Lett. 2014 Jul; 113:028101. https://link.aps.org/doi/10.1103/PhysRevLett.113.028101, doi: 10.1103/Phys-RevLett.113.028101.

Iyer-Biswas S, Wright CS, Henry JT, Lo K, Burov S, Lin Y, Crooks GE, Crosson S, Dinner AR, Scherer NF. Scaling laws governing stochastic growth and division of single bacterial cells. Proc Natl Acad Sci USA. 2014; 111(45):15912–15917. doi: 10.1073/pnas.1403232111.

Jia C, Grima R. Frequency Domain Analysis of Fluctuations of mRNA and Protein Copy Numbers within a Cell Lineage: Theory and Experimental Validation. Phys Rev X. 2021 May; 11:021032. https://link.aps.org/doi/10.1103/PhysRevX.11.021032, doi: 10.1103/PhysRevX.11.021032.

Jia C, Singh A, Grima R. Concentration fluctuations in growing and dividing cells: Insights into the emergence of concentration homeostasis. PLoS Comput Biol. 2022 10; 18(10):1–34. https://doi.org/10.1371/journal.pcbi.1010574, doi: 10.1371/journal.pcbi.1010574.

Johnston IG, Jones NS. Closed-form stochastic solutions for non-equilibrium dynamics and inheritance of cellular components over many cell divisions. Proc R Soc A: Math Phys Eng Sci. 2015; 471(2180):20150050. https://royalsocietypublishing.org/doi/abs/10.1098/rspa.2015.0050, doi: 10.1098/rspa.2015.0050.

Jedrak J, Kwiatkowski M, Ochab-Marcinek A. Exactly solvable model of gene expression in a proliferating bacterial cell population with stochastic protein bursts and protein partitioning. Phys Rev E. 2019 Apr; 99:042416. https://link.aps.org/doi/10.1103/PhysRevE.99.042416, doi: 10.1103/PhysRevE.99.042416.

Krächan EG, Fischer AU, Franke J, Friauf E. Synaptic reliability and temporal precision are achieved via high quantal content and effective replenishment: auditory brainstem versus hippocampus. J Physiol. 2017; 595(3):839–864. doi: 10.1113/JP272799.

Little JW, Michalowski CB. Stability and Instability in the Lysogenic State of Phage Lambda. J Bacteriol. 2010; 192(22):6064–6076. https://journals.asm.org/doi/abs/10.1128/jb.00726-10, doi: 10.1128/jb.00726-10.

Liu X, Yan J, Kirschner MW. Cell size homeostasis is tightly controlled throughout the cell cycle. PLoS Biol. 2024; 22(1):e3002453. doi: 10.1371/journal.pbio.3002453.

Losick R, Desplan C. Stochasticity and Cell Fate. Science. 2008; 320(5872):65–68. https://www.science.org/doi/abs/10.1126/science.1147888, doi: 10.1126/science.1147888.

Lovatt D, Ruble BK, Lee, et al. Transcriptome in vivo analysis (TIVA) of spatially defined single cells in live tissue. Nat Methods. 2014 Feb; 11(2):190–196. https://doi.org/10.1038/nmeth.2804, doi: 10.1038/nmeth.2804.

Luo L, Bai Y, Fu X. Stochastic threshold in cell size control. Phys Rev Res. 2023 Mar; 5:013173. https://link.aps.org/doi/10.1103/PhysRevResearch.5.013173, doi: 10.1103/PhysRevResearch.5.013173.

Mahdessian D, Cesnik AJ, Gnann C, et al. Spatiotemporal dissection of the cell cycle with single-cell proteogenomics. Nature. 2021; 590(7847):649–654. doi: 10.1038/s41586-021-03232-9.

Männik J, Kar P, Amarasinghe C, Amir A, Männik J. Determining the rate-limiting processes for cell division in Escherichia coli. Nat Commun. 2024; 15(1):9948. doi: 10.1038/s41467-024-54242-w.

Miyazaki R, Minoia M, Pradervand N, Sulser S, Reinhard F, van der Meer JR. Cellular Variability of RpoS Expression Underlies Subpopulation Activation of an Integrative and Conjugative Element. PLoS Genet. 2012 07; 8(7):1–15. https://doi.org/10.1371/journal.pgen.1002818, doi: 10.1371/journal.pgen.1002818.

Munsky B, Neuert G, van Oudenaarden A. Using Gene Expression Noise to Understand Gene Regulation. Science. 2012; 336(6078):183–187. https://www.science.org/doi/abs/10.1126/science.1216379, doi: 10.1126/science.1216379.

Nieto C, Augusto Vargas-Garcia C, Singh A. A Moments-Based Analytical Approach for Cell Size Homeostasis. IEEE Control Syst Lett. 2024; 8:2205–2210. doi: 10.1109/LCSYS.2024.3411041.

Nieto C, Vargas-García CA, Pedraza JM, Singh A. Mechanisms of cell size regulation in slow-growing escherichia coli cells: Discriminating models beyond the adder. npj Syst Biol Appl. 2024; 10(1):61. doi: 10.1038/s41540024-00383-z.

Nieto C, Vargas-García CA, Singh A. A generalized adder for cell size homeostasis: Effects on stochastic clonal proliferation. Biophys J. 2025; 124(9):1376–1386. https://www.sciencedirect.com/science/article/pii/S0006349525001651, doi: 10.1016/j.bpj.2025.03.011.

Padovan-Merhar O, Nair GP, Biaesch AG, et al. Single Mammalian Cells Compensate for Differences in Cellular Volume and DNA Copy Number through Independent Global Transcriptional Mechanisms. Mol Cell. 2015; 58(2):339–352. doi: 10.1016/j.molcel.2015.03.005.

Padovan-Merhar O, Raj A. Using variability in gene expression as a tool for studying gene regulation. WIREs Syst Biol Med. 2013; 5(6):751–759. https://wires.onlinelibrary.wiley.com/doi/abs/10.1002/wsbm.1243, doi: 10.1002/wsbm.1243.

Paulsson J. Models of stochastic gene expression. Phys Life Rev. 2005; 2(2):157–175. https://www.sciencedirect.com/science/article/pii/S1571064505000138, doi: 10.1016/j.plrev.2005.03.003.

Perez-Carrasco R, Beentjes C, Grima R. Effects of cell cycle variability on lineage and population measurements of messenger RNA abundance. J R Soc Interface. 2020; 17(168):20200360. doi: 10.1098/rsif.2020.0360.

Powell EO. Growth Rate and Generation Time of Bacteria, with Special Reference to Continuous Culture. Microbiol. 1956; 15(3):492–511. https://www.microbiologyresearch.org/content/journal/micro/10.1099/00221287-15-3-492, doi: 10.1099/00221287-15-3-492.

Raj A, van den Bogaard P, Rifkin SA, van Oudenaarden A, Tyagi S. Imaging individual mRNA molecules using multiple singly labeled probes. Nat Methods. 2008 Oct; 5(10):877–879. https://doi.org/10.1038/nmeth.1253, doi: 10.1038/nmeth.1253.

Raj A, Peskin CS, Tranchina D, Vargas DY, Tyagi S. Stochastic mRNA Synthesis in Mammalian Cells. PLoS Biol. 2006; 4(10):e309. https://doi.org/10.1371/journal.pbio.0040309.

Raj A, Van Oudenaarden A. Nature, Nurture, or Chance: Stochastic Gene Expression and Its Consequences. Cell. 2008; 135(2):216–226. doi: 10.1016/j.cell.2008.09.050.

Raser JM, O’Shea EK. Control of Stochasticity in Eukaryotic Gene Expression. Science. 2004; 304(5678):1811– 1814. https://www.science.org/doi/abs/10.1126/science.1098641, doi: 10.1126/science.1098641.

Reshes G, Vanounou S, Fishov I, Feingold M. Timing the start of division in E. coli: a single-cell study. Phys Biol. 2008 nov; 5(4):046001. https://dx.doi.org/10.1088/1478-3975/5/4/046001, doi: 10.1088/1478-3975/5/4/046001.

Reshes G, Vanounou S, Fishov I, Feingold M. Cell Shape Dynamics in Escherichia coli. Biophy J. 2008; 94(1):251– 264. doi: 10.1529/biophysj.107.104398.

Rigney DR. Stochastic model of constitutive protein levels in growing and dividing bacterial cells. J Theor Biol. 1979; 76(4):453–480. https://www.sciencedirect.com/science/article/pii/0022519379900134, doi: 10.1016/0022-5193(79)90013-4.

Rijal K, Müller NIC, Friauf E, Singh A, Prasad A, Das D. Exact Distribution of the Quantal Content in Synaptic Transmission. Phys Rev Lett. 2024 May; 132:228401. https://link.aps.org/doi/10.1103/PhysRevLett.132.228401, doi: 10.1103/PhysRevLett.132.228401.

Rijal K, Prasad A, Singh A, Das D. Exact Distribution of Threshold Crossing Times for Protein Concentrations: Implication for Biological Timekeeping. Phys Rev Lett. 2022 Jan; 128:048101. https://link.aps.org/doi/10.1103/PhysRevLett.128.048101, doi: 10.1103/PhysRevLett.128.048101.

Roeder AHK, Chickarmane V, Cunha A, Obara B, Manjunath BS, Meyerowitz EM. Variability in the Control of Cell Division Underlies Sepal Epidermal Patterning in Arabidopsis thaliana. PLoS Biol. 2010 05; 8(5):1–17. doi: 10.1371/journal.pbio.1000367.

Saint-Antoine MM, Singh A. Network inference in systems biology: recent developments, challenges, and applications. Curr Opin Biotechnol. 2020; 63:89–98. https://www.sciencedirect.com/science/article/pii/S0958166919301399, doi: 10.1016/j.copbio.2019.12.002.

Sanchez A, Garcia HG, Jones D, Phillips R, Kondev J. Effect of Promoter Architecture on the Cell-to-Cell Variability in Gene Expression. PLoS Comp Biol. 2011 03; 7(3):1–20. https://doi.org/10.1371/journal.pcbi.1001100, doi: 10.1371/journal.pcbi.1001100.

Sanchez A, Golding I. Genetic Determinants and Cellular Constraints in Noisy Gene Expression. Science. 2013; 342(6163):1188–1193. https://www.science.org/doi/abs/10.1126/science.1242975, doi: 10.1126/science.1242975.

Schuh L, Saint-Antoine M, Sanford EM, et al. Gene Networks with Transcriptional Bursting Recapitulate Rare Transient Coordinated High Expression States in Cancer. Cell Syst. 2020; 10(4):363–378.e12. https://doi.org/10.1016/j.cels.2020.03.004, doi: 10.1016/j.cels.2020.03.004.

Shaffer SM, Dunagin MC, Torborg SR, et al. Rare cell variability and drug-induced reprogramming as a mode of cancer drug resistance. Nature. 2017; 546(7658):431–435. doi: 10.1038/nature22794.

Shaffer SM, Emert BL, Reyes Hueros RA, et al. Memory Sequencing Reveals Heritable Single-Cell Gene Expression Programs Associated with Distinct Cellular Behaviors. Cell. 2020; 182(4):947–959.e17. https://doi.org/10.1016/j.cell.2020.07.003, doi: 10.1016/j.cell.2020.07.003.

Shahrezaei V, Swain PS. Analytical distributions for stochastic gene expression. Proc Natl Acad Sci USA. 2008; 105(45):17256–17261. https://www.pnas.org/doi/abs/10.1073/pnas.0803850105.

Singh A, Razooky BS, Dar RD, Weinberger LS. Dynamics of protein noise can distinguish between alternate sources of gene-expression variability. Mol Syst Biol. 2012; 8(1):607. doi: 10.1038/msb.2012.38.

Singh A, Vargas CA, Karmakar R. Stochastic analysis and inference of a two-state genetic promoter model. In: 2013 American Control Conference IEEE; 2013. p. 4563–4568. doi: 10.1109/ACC.2013.6580542.

Singh A, Weinberger LS. Stochastic gene expression as a molecular switch for viral latency. Curr Opin Microbiol. 2009; 12(4):460–466. doi: 10.1016/j.mib.2009.06.016.

Soltani M, Singh A. Effects of cell-cycle-dependent expression on random fluctuations in protein levels. R Soc Open Sci. 2016; 3(12):160578. https://royalsocietypublishing.org/doi/abs/10.1098/rsos.160578, doi: 10.1098/rsos.160578.

Soltani M, Vargas-Garcia CA, Antunes D, Singh A. Intercellular Variability in Protein Levels from Stochastic Expression and Noisy Cell Cycle Processes. PLoS Comput Biol. 2016 08; 12:1–23. https://doi.org/10.1371/journal.pcbi.1004972, doi: 10.1371/journal.pcbi.1004972.

Storz G, Opdyke JA, Zhang A. Controlling mRNA stability and translation with small, noncoding RNAs. Curr Opin Microbiol. 2004; 7(2):140–144. doi: 10.1016/j.mib.2004.02.015.

Stukalin EB, Aifuwa I, Kim JS, Wirtz D, Sun SX. Age-dependent stochastic models for understanding population fluctuations in continuously cultured cells. J R Soc Interface. 2013; 10(85):20130325. https://royalsocietypublishing.org/doi/abs/10.1098/rsif.2013.0325, doi: 10.1098/rsif.2013.0325.

Suen TK, A. B, Placek K. Cell-to-cell proteome variability: life in a cycle. Sig Transduct Target Ther. 2021; 6(1):229. doi: 10.1038/s41392-021-00655-8.

Swain PS, Elowitz MB, Siggia ED. Intrinsic and extrinsic contributions to stochasticity in gene expression. Proc Natl Acad Sci USA. 2002; 99(20):12795–12800. https://www.pnas.org/doi/abs/10.1073/pnas.162041399, doi: 10.1073/pnas.162041399.

Taheri-Araghi S, Bradde S, Sauls JT, Hill NS, Levin PA, Paulsson J, Vergassola M, Jun S. Cell-Size Control and Homeostasis in Bacteria. Curr Bio. 2015; 25(3):385–391. doi: 10.1016/j.cub.2014.12.009.

Taniguchi Y, Choi PJ, Li GW, Chen H, Babu M, Hearn J, Emili A, Xie XS. Quantifying E. coli Proteome and Transcriptome with Single-Molecule Sensitivity in Single Cells. Science. 2010; 329(5991):533–538. https://www.science.org/doi/abs/10.1126/science.1188308, doi: 10.1126/science.1188308.

Thattai M, Van Oudenaarden A. Intrinsic noise in gene regulatory networks. Proc Natl Acad Sci USA. 2001; 98(15):8614–8619. https://doi.org/10.1073/pnas.151588598.

Thomas P. Intrinsic and extrinsic noise of gene expression in lineage trees. Sci Rep. 2019; 9(1):474. doi: 10.1038/s41598-018-35927-x.

Tsukanov R, Reshes G, Carmon G, Fischer-Friedrich E, Gov NS, Fishov I, Feingold M. Timing of Z-ring localization in Escherichia coli. Phys Biol. 2011 oct; 8(6):066003. https://dx.doi.org/10.1088/1478-3975/8/6/066003, doi: 10.1088/1478-3975/8/6/066003.

Vahdat Z, Gambrell O, Fisch J, Friauf E, Singh A. Inferring synaptic transmission from the stochastic dynamics of the quantal content: An analytical approach. PLoS Comput Biol. 2025; 21(5):e1013067. doi: 10.1371/journal.pcbi.1013067.

Vargas-Garcia CA, Ghusinga KR, Singh A. Cell size control and gene expression homeostasis in single-cells. Curr Opin Syst Biol. 2018; 8:109–116. doi: 10.1016/j.coisb.2018.01.002.

Wang Y, Yu Z, Grima R, Cao Z. Exact solution of a three-stage model of stochastic gene expression including cell-cycle dynamics. J Chem Phys. 2023 12; 159(22):224102. https://doi.org/10.1063/5.0173742, doi: 10.1063/5.0173742.

Yates CA, Ford MJ, Mort RL. A Multi-stage Representation of Cell Proliferation as a Markov Process. Bull Math Biol. 2017; 79:2905–2928. doi: 10.1007/s11538-017-0356-4.

Yu J, Xiao J, Ren X, Lao K, Xie XS. Probing Gene Expression in Live Cells, One Protein Molecule at a Time. Science. 2006; 311(5767):1600–1603. https://www.science.org/doi/abs/10.1126/science.1119623, doi: 10.1126/science.1119623.

Zhang Z, Zabaikina I, Nieto C, Vahdat Z, Bokes P, Singh A. Stochastic Gene Expression in Proliferating Cells: Differing Noise Intensity in Single-Cell and Population Perspectives. bioRxiv. 2024; doi: 10.1101/2024.06.28.601263.

Zilman A, Ganusov VV, Perelson AS. Stochastic Models of Lymphocyte Proliferation and Death. PLoS One. 2010 09; 5(9):1–14. doi: 10.1371/journal.pone.0012775.

Zopf CJ, Quinn K, Zeidman J, Maheshri N. Cell-Cycle Dependence of Transcription Dominates Noise in Gene Expression. PLoS Comput Biol. 2013 07; 9(7):1–12. https://doi.org/10.1371/journal.pcbi.1003161, doi: 10.1371/journal.pcbi.1003161.

