## Supplementary material for "Cyclo-stationary distributions of mRNA and Protein counts for random cell division times": Manuscript

### I. GENERAL FRAMEWORK TO STUDY THE CYCLO-STATIONARY DISTRIBUTIONS

#### A. Equation for the cyclo-stationary copy number distributions $P_+^{ss}(y_+)$ and $P_-^{ss}(y_-)$

In the main text, we present the equations for $P_+^i(y_{+,i}, t_{s_i})$ and $P_-^i(y_{-,i}, t_{s_i})$, the distributions of copy numbers just after and just before the $i^{th}$ cell division respectively, as follows:

$$P_+^i(y_{+,i}, t_{s_i}) = \sum_{y_{+,i-1}} \sum_{x_{+,i}} B(y_{+,i} + x_{+,i}, \frac{1}{2}, x_{+,i}) p(y_{+,i} + x_{+,i}, t_{s_i} | y_{+,i-1}) \; P_+^{i-1}(y_{+,i-1}, t_{s_{i-1}}), \tag{1}$$

$$P_-^i(y_{-,i}, t_{s_i}) = \sum_{y_{+,i-1}} p(y_{-,i}, t_{s_i} | y_{+,i-1}) \; P_+^{i-1}(y_{+,i-1}, t_{s_{i-1}}). \tag{2}$$

Integrating Eq. 1 over the joint probability distribution of $g_2(t_{s_i}, t_{s_{i-1}})$ of successive division time intervals,

$$\int_0^\infty \int_0^\infty dt_{s_i} dt_{s_{i-1}} g_2(t_{s_i}, t_{s_{i-1}}) P_+^i(y_{+,i}, t_{s_i}) = \int_0^\infty \int_0^\infty dt_{s_i} dt_{s_{i-1}} g_2(t_{s_i}, t_{s_{i-1}}) \sum_{y_{+,i-1}} \sum_{x_{+,i}} B(y_{+,i} + x_{+,i}, \frac{1}{2}, x_{+,i}) \\ p(y_{+,i} + x_{+,i} | y_{+,i-1}, t_{s_i}) P_+^i(y_{+,i-1}, t_{s_{i-1}}). \tag{3}$$

If it is further assumed that successive division times are uncorrelated, i.e. $g_2(t_{s_i}, t_{s_{i-1}}) = g(t_{s_i}) g(t_{s_{i-1}})$, where $g(t_{s_i})$ is the normalized distributions of $t_{s_i}$, we have

$$\int_0^\infty \int_0^\infty dt_{s_i} dt_{s_{i-1}} g(t_{s_i}) g(t_{s_{i-1}}) P_+^i(y_{+,i}, t_{s_i}) = \int_0^\infty \int_0^\infty dt_{s_i} dt_{s_{i-1}} g(t_{s_i}) g(t_{s_{i-1}}) \sum_{y_{+,i-1}} \sum_{x_{+,i}} B(y_{+,i} + x_{+,i}, \frac{1}{2}, x_{+,i}) \\ p(y_{+,i} + x_{+,i} | y_{+,i-1}, t_{s_i}) P_+^i(y_{+,i-1}, t_{s_{i-1}}) \tag{4}$$

which simplifies to

$$\int_0^\infty dt_{s_i} g(t_{s_i}) P_+^i(y_{+,i}, t_{s_i}) = \sum_{y_{+,i-1}, x_{+,i}} \int_0^\infty dt_{s_i} g(t_{s_i}) p(y_{+,i} + x_{+,i} | y_{+,i-1}, t_{s_i}) \; B\left(y_{+,i} + x_{+,i}, \frac{1}{2}, x_{+,i}\right) \\ \times \int_0^\infty dt_{s_{i-1}} g(t_{s_{i-1}}) P_+^i(y_{+,i-1}, t_{s_{i-1}}) \tag{5}$$

For $i \gg 1$, as the cyclo-stationary regime is attained, we may define the distribution as cell birth, $P_+^{ss}(y_+) = \int_0^\infty dt_{s_i} g(t_{s_i}) P_+^i(y_{+,i}, t_{s_i})$. Dropping the subscripts $i$, and setting $t_{s_i} = t_s$ and $y_{+,i-1} = y'_+$, Eq. 5 gives

$$P_+^{ss}(y_+) = \sum_{y'_+} \sum_{x_+} \int_0^\infty dt_s g(t_s) \; B(y_+ + x_+, \frac{1}{2}, x_+) \times p(y_+ + x_+, t_s | y'_+) \; P_+^{ss}(y'_+). \tag{6}$$

In a similar way, one may derive from Eq. 2 above, the cyclo-stationary distribution just before division, defined as $P_-^{ss}(y_-) = \int_0^\infty dt_{s_i} g(t_{s_i}) P_-^i(y_{-,i}, t_{s_i})$, related to $P_+^{ss}(y_+)$:

$$P_-^{ss}(y_-) = \sum_{y'_+} \int_0^\infty dt_s g(t_s) \; p(y_-, t_s | y'_+, t_s) \; P_+^{ss}(y'_+). \tag{7}$$

Eqs. 6 and 7 are the two Eqs. 3 and 4 in the main text.

#### B. Self-consistent integral for the generating functions $F_+(q)$ and its relation to $F_-(q)$

We define generating function $F_\pm(q) = \sum_{y_\pm=0}^\infty q^{y_\pm} P_\pm^{ss}(y_\pm)$. Multiply $\sum_{y_+} q^{y_+}$ on both sides of Eq. 6 we get,

$$F_+(q) = \sum_{y'_+} \sum_{x_+} \int_0^\infty dt_s g(t_s) \sum_{y_+} q^{y_+} p(y_+ + x_+, t_s | y'_+) \; B(y_+ + x_+, \frac{1}{2}, x_+) P_+^{ss}(y'_+) \tag{8}$$

We defining a new variable $\tilde{y}_+ = y_+ + x_+$, where $x_+ \le \tilde{y}_+ \le \infty$. But since $B(\tilde{y}_+, \frac{1}{2}, x_+) = 0$ for $\tilde{y}_+ < x_+$, we put $\tilde{y}_+ \in [0, \infty)$. Using the explicit form of the binomial function, the recursion relation of $F_+(q)$ may then be written as:

$$\begin{aligned} F_+(q) &= \sum_{y'_+} \int_0^\infty dt_s g(t_s) \sum_{\tilde{y}_+} q^{\tilde{y}_+} p(\tilde{y}_+, t_s|y'_+) \sum_{x_+=0}^{\tilde{y}_+} \frac{1}{q^{x_+}} \binom{\tilde{y}_+}{x_+} \left(\frac{1}{2}\right)^{\tilde{y}_+ - x_+} \left(\frac{1}{2}\right)^{x_+} P_+^{ss}(y'_+) \\ &= \sum_{y'_+} \int_0^\infty dt_s g(t_s) \left( \sum_{\tilde{y}_+} q^{\tilde{y}_+} p(\tilde{y}_+, t_s|y'_+) \left(\frac{q+1}{2q}\right)^{\tilde{y}_+} \right) P_+^{ss}(y'_+) \\ &= \int_0^\infty dt_s g(t_s) \sum_{y'_+} F\left(\frac{q+1}{2}, t_s \Big| y'_+\right) P_+^{ss}(y'_+). \end{aligned} \tag{9}$$

Here $F(q, t|y') = \sum_y q^y p(y, t|y')$ is the generating function of the probability $p(y, t|y')$ tied to the process of gene expression. Let us assume that this generating function has a form $F(q, t|y') = \mathcal{H}(q-1, \gamma_y t) \times (1 + (q-1)e^{-\gamma_y t})^{y'}$. This leads to the following self-consistent integral for $F_+(q)$ (which is Eq. 6 in the main text):

$$F_+(q) = \int_0^\infty dt_s g(t_s) \mathcal{H}(\frac{q-1}{2}, \gamma_y t_s) F_+ \left(1 + \frac{(q-1)}{2} e^{-\gamma_y t_s}\right). \tag{10}$$

In a similar way, by multiplying $\sum_{y_-} q^{y_-}$ on both sides of Eq. 7 we get,

$$\begin{aligned} F_-(q) &= \sum_{y_-} q^{y_-} \sum_{y'_+} \int_0^\infty dt_s g(t_s) \times p(y_-, t_s|y'_+) \times P_+^{ss}(y'_+) \\ &= \int_0^\infty dt_s g(t_s) \sum_{y'_+} F(q, t_s|y'_+) P_+^{ss}(y'_+) \\ &= \int_0^\infty dt_s g(t_s) \mathcal{H}(q-1, \gamma_y t_s) F_+ \left(1 + (q-1)e^{-\gamma_y t_s}\right). \end{aligned} \tag{11}$$

Putting $q = 2q' - 1$ in Eq. 10 we have,

$$F_+(2q'-1) = \int_0^\infty dt_s g(t_s) \mathcal{H}(q'-1, \gamma_y t_s) F_+ \left(1 + (q'-1)e^{-\gamma_y t_s}\right), \tag{12}$$

and comparing with Eq. 11, we obtain (the Eq. 7 in the main text):

$$F_-(q) = F_+(2q-1) \tag{13}$$

##### C. Closed form of $F_+(q)$ for Fixed cell cycle times, and the intractable nested integrals for Random cell cycle times

For fixed cell division times, i.e. $g(t_s) = \delta(t_s - T)$, Eq. 10 reduces to

$$F_+(q) = \mathcal{H}\left(\frac{q-1}{2}, \gamma_y T\right) F_+ \left(1 + \frac{(q-1)}{2} e^{-\gamma_y T}\right). \tag{14}$$

If we set, $q - 1 = w$ then Eq. 14 becomes to

$$F_+(1+w) = \mathcal{H}\left(\frac{w}{2}, \gamma_y T\right) F_+ \left(1 + \frac{w}{2} e^{-\gamma_y T}\right). \tag{15}$$

This recursion formula may be iterated to obtain

$$F_+(1+w) = \prod_{k=1}^{j} \mathcal{H}\left(\frac{w}{2} \frac{e^{-(k-1)\gamma_y T}}{2^{k-1}}, \gamma_y T\right) F_+ \left(1 + w\left(\frac{e^{-\gamma_y T}}{2}\right)^j\right). \tag{16}$$

As $j \to \infty$, $F_+\left(1 + w\left(\frac{e^{-\gamma_y T}}{2}\right)^j\right) \to F_+(1) = 1$, and hence

$$F_+(1+w) = \prod_{k=1}^{\infty} \mathcal{H}\left(\frac{w}{2}\frac{e^{-(k-1)\gamma_y T}}{2^{k-1}}, \gamma_y T\right). \tag{17}$$

Replacing back $w = q - 1$ Eq. 17 gives the closed form in Eq. 8 in the main text.

*For random division times* $t_s$, with any general normalised function $g(t_s)$, when we substitute $w = q - 1$, Eq. 10 becomes:

$$F_+(1+w) = \int_0^\infty dt_s\ g(t_s)\mathcal{H}\left(\frac{w}{2}, \gamma_y t_s\right) F_+\left(1 + \frac{w}{2}e^{-\gamma_y t_s}\right). \tag{18}$$

Iterating one step, and replacing the $F_+$ on the right side with an similar integral as Eq. 18, we obtain:

$$F_+(w+1) = \int_0^\infty dt_s^1\ g(t_s^1)\mathcal{H}\left(\frac{w}{2}, \gamma_y t_s^1\right) \int_0^\infty dt_s^2\ g(t_s^2)\mathcal{H}\left(\frac{w_1(t_s^1)}{2}, \gamma_y t_s^2\right) F_+\left(1 + \frac{w_1(t_s^1)}{2}e^{-\gamma_y t_s^2}\right), \tag{19}$$

where $w_1(t_s^1) = \frac{w}{2}e^{-\gamma_y t_s^1}$. Continuing with the next iteration,

$$F_+(w+1) = \int_0^\infty dt_s^1\ g(t_s^1)\mathcal{H}\left(\frac{w}{2}, \gamma_y t_s^1\right) \int_0^\infty dt_s^2\ g(t_s^2)\mathcal{H}\left(\frac{w_1}{2}, \gamma_y t_s^2\right) \int_0^\infty dt_s^3\ g(t_s^3)\mathcal{H}\left(\frac{w_2}{2}, \gamma_y t_s^3\right) F_+\left(1 + \frac{w_2}{2}e^{-\gamma_y t_s^3}\right), \tag{20}$$

where $w_2 = w_2(t_s^1, t_s^2) = \frac{w_1}{2}e^{-\gamma_y t_s^2} = \frac{w}{2^2}e^{-\gamma_y t_s^1}e^{-\gamma_y t_s^2}$. Repeating this indefinitely, as $j \to \infty$ we have $F_+(1 + w_j) \to F_+(1) = 1$, where $w_j = \frac{w_{j-1}}{2}e^{-\gamma_y t_s^j}$, and hence

$$F_+(1+w) = \prod_{k=1}^{\infty} \int_0^\infty dt_s^k\ g(t_s^k)\mathcal{H}\left(\frac{w_{k-1}}{2}, \gamma_y t_s^k\right). \tag{21}$$

As $w_k = w_k(t_s^1, t_s^2, ..., t_s^k) = \frac{w}{2^k}e^{-\gamma_y t_s^1}e^{-\gamma_y t_s^2}...e^{-\gamma_y t_s^k}$, the above nested integrals are in general intractable. This is why the problem has stayed challenging.

##### D. Deriving the cyclo-stationary distributions at birth and before division, from the series expansion of generating function about $q = 1$

Although $P_+^{ss}(y_+)$ are themselves coefficients of the series expansion of $F_+(q)$ about $q = 0$, we may start with an alternate expansion of $F_+(q) = \sum_{k=0}^{\infty} \frac{(q-1)^k}{k!} F_+^{(k)}(1)$ about $q = 1$. In that case,

$$P_+^{ss}(y_+) = \frac{1}{y_+!}\left[\frac{\partial^{y_+}}{\partial q^{y_+}} F_+(q)\right]_{q=0} = \frac{1}{y_+!}\left[\frac{\partial^{y_+}}{\partial q^{y_+}} \sum_{k=0}^{\infty} \frac{(q-1)^k}{k!} F_+^{(k)}(1)\right]_{q=0} = \sum_{k=y_+}^{\infty} \binom{k}{y_+} \frac{(-1)^{k-y_+}}{k!} F_+^{(k)}(1) \tag{22}$$

Similarly, using Eq. 13, we have

$$\begin{aligned} P_-^{ss}(y_-) = \frac{1}{y_-!}\left[\frac{\partial^{y_-}}{\partial q^{y_-}} F_-(q)\right]_{q=0} &= \frac{1}{y_-!}\left[\frac{\partial^{y_-}}{\partial q^{y_-}} F_+(2q-1)\right]_{q=0} = \frac{1}{y_-!}\left[\frac{\partial^{y_-}}{\partial q^{y_-}} \sum_{k=0}^{\infty} \frac{2^k (q-1)^k}{k!} F_+^{(k)}(1)\Big|_{q=0}\right] \\ &= \sum_{k=y_-}^{\infty} \binom{k}{y_-} \frac{(-1)^{k-y_-}}{k!} 2^k F_+^{(k)}(1) \end{aligned} \tag{23}$$

Thus above, we have the series expansions of $P_+^{ss}(y_+)$ and $P_-^{ss}(y_-)$ (Eq. 11 and 12 in the main text) involving the coefficients $F_+^{(k)}(1)$.

##### E. The first three cumulants of $P_+^{ss}(y_+)$ in terms of the coefficients $F_+^{(k)}(1)$

Using $F_+(q) = \sum_{y_+} P_+^{ss}(y_+) q^{y_+} = \sum_{k=0}^{\infty} \frac{(q-1)^k}{k!} F_+^{(k)}(1)$, we may obtain the cumulants as follows. The mean of $y_+$:

$$\langle y_+ \rangle = \sum_{y_+} y_+ P_+^{ss}(y_+) = q\frac{\partial}{\partial q} F_+(q)\Big|_{q=1} = q\frac{\partial}{\partial q}\left(\sum_k \frac{(q-1)^k}{k!} F_+^{(k)}(1)\right)_{q=1} = q\sum_k \frac{k(q-1)^{k-1}}{k!} F_+^{(k)}(1)\Big|_{q=1} = F_+^{(1)}(1) \tag{24}$$

The second moment

$$\langle y_+^2 \rangle = q\frac{\partial}{\partial q} q\frac{\partial}{\partial q} F_+(q)\Big|_{q=1} = q\sum_k \frac{k(q-1)^{k-1}}{k!} F_+^{(k)}(1) + q^2 \sum_k \frac{k(k-1)(q-1)^{k-2}}{k!} F_+^{(k)}(1)\Big|_{q=1} = F_+^{(1)}(1) + F_+^{(2)}(1) \tag{25}$$

Hence the Variance

$$\kappa_2 = \langle y_+^2 \rangle - \langle y_+ \rangle^2 = F_+^{(1)}(1) + F_+^{(2)}(1) - \left(F_+^{(1)}(1)\right)^2 \tag{26}$$

The third moment

$$\begin{aligned}\langle y_+^3 \rangle &= q\frac{d}{dq} q\frac{d}{dq} q\frac{d}{dq} F_+(q)\Big|_{q=1} \\ &= q\sum_k \frac{k(q-1)^{k-1}}{k!} F_+^{(k)}(1) + 3q^2 \sum_k \frac{k(k-1)(q-1)^{k-2}}{k!} F_+^{(k)}(1) + q^2 \sum_k \frac{k(k-1)(k-2)(q-1)^{k-3}}{k!} F_+^{(k)}(1)\Big|_{q=1} \\ &= F_+^{(1)}(1) + 3F_+^{(2)}(1) + F_+^{(3)}(1)\end{aligned} \tag{27}$$

Hence the third cumulant is (see Eqs. 24, 26 and 27 above)

$$\kappa_3 = \langle (y_+ - \kappa_1)^3 \rangle = [F_+^{(1)}(1) + 3F_+^{(2)}(1) + F_+^{(3)}(1)] - 3\kappa_1\kappa_2 - \kappa_1^3 \tag{28}$$

The above equations appear as Eq. 13, 14 and 15 in the main text. Using the above cumulants we obtain $CV^2 = \kappa_2/\kappa_1^2$ and Skewness $= \kappa_3/\kappa_2^{3/2}$ in our study.

#### II. STATISTICS OF THE mRNA NUMBER IN THE CYCLO-STATIONARY STATE

##### A. The generating function related to the model of transcription, and thereby obtaining function $\mathcal{H}$

The Master equation for the stochastic model of mRNA production and degradation is

$$\frac{dp(m,t|m'_+)}{dt} = k_m p(m-1,t|m'_+) + \gamma_m (m+1) p(m+1,t|m'_+) - (\gamma_m m + k_m) p(m,t|m'_+). \tag{29}$$

Here $k_m$ is the transcription rate, and $\gamma_m$ is the degradation rate of mRNAs. The generating function $F(q,t) = \sum_{j=0}^{\infty} q^m p(m,t|m'_+)$ of the distribution $p(m,t|m'_+)$ satisfies (using Eq. 29 above) the following:

$$\frac{\partial F(q,t)}{\partial t} + \gamma_m (q-1)\frac{\partial F}{\partial q} = k_m (q-1) F. \tag{30}$$

Eq. 30 can be solved by using the method of Lagrange characteristic, and one gets

$$F(q,t) = e^{\lambda(t)(q-1)}(1 + (q-1)e^{-\gamma_m t})^{m'_+}, \tag{31}$$

where $\lambda(t) = (k_m/\gamma_m)[1 - e^{-\gamma_m t}]$. For brevity we will use $\lambda(t) \equiv \lambda$ below. Thus comparing with Eq. 5 of the main text (also see below Eq. 9), we identify the function

$$\mathcal{H} = e^{\frac{k_m}{\gamma_m}[1-e^{-\gamma_m t}](q-1)}. \tag{32}$$

##### B. Obtaining the coefficients $F_+^{(k)}(1)$ and the series of the distributions $P_\pm^{ss}(m_\pm)$

Using $\mathcal{H}$ from Eq. 32 in Eq. 10, and $F_+(q) = \sum_{j=0}^{\infty} \frac{(q-1)^j}{j!} F_+^{(j)}(1)$ we have

$$\begin{aligned}
F_+(q) &= \int_0^\infty dt_s g(t_s) e^{\lambda((q-1)/2)} F_+(1 + ((q-1)/2)e^{-\gamma_m t_s}) \\
&= \int_0^\infty dt_s g(t_s) e^{\lambda((q-1)/2)} \sum_{j=0}^{\infty} \frac{F_+^{(j)}(1)}{j!} \left(\frac{q-1}{2}\right)^j e^{-j\gamma_m t_s} \\
&= \int_0^\infty dt_s g(t_s) \sum_{l=0}^{\infty}\sum_{j=0}^{\infty} \frac{\lambda^l \left(\frac{q-1}{2}\right)^l}{l!} \frac{F_+^{(j)}(1)}{j!} \left(\frac{q-1}{2}\right)^j e^{-j\gamma_m t_s} \\
&= \int_0^\infty dt_s g(t_s) \sum_{l=0}^{\infty}\sum_{j=0}^{\infty} \frac{F_+^{(j)}(1)}{l!j!} \left(\frac{q-1}{2}\right)^{l+j} \left(\frac{k_m}{\gamma_m}\right)^l e^{-j\gamma_m t_s}(1-e^{-\gamma_m t_s})^l
\end{aligned} \tag{33}$$

Changing summation indices to $k = l + j$ and defining $\Psi_{k,j} = \int_0^\infty dt_s g(t_s) e^{-j\gamma_m t_s}(1-e^{-\gamma_m t_s})^{k-j}$, Eq. 33 becomes

$$F_+(q) = \sum_{k=0}^{\infty} \frac{1}{k!}\left(\frac{q-1}{2}\right)^k \sum_{j=0}^{k} \binom{k}{j} \left(\frac{k_m}{\gamma_m}\right)^{k-j} \Psi_{k,j} F_+^{(j)}(1) \tag{34}$$

Using the relation $F_+(q) = \sum_{k=0}^{\infty} \frac{(q-1)^k}{k!} F_+^k(1)$ on the left side of Eq. 34 above, and comparing coefficients we get the desired recursion relation (which appears in Eq. 21 of the main text):

$$F_+^k(1) = \frac{1}{2^k} \sum_{j=0}^{k} \left(\frac{k_m}{\gamma_m}\right)^{k-j} \binom{k}{j} \Psi_{k,j} F_+^j(1). \tag{35}$$

The first few coefficients are explicitly as follows. As $\sum_{m_+} P^{ss}(m_+) = 1$ we firstly have $F_+^{(0)}(1) = 1$. The next coefficient (from Eq. 35) is

$$F_+^1(1) = \frac{1}{2}\left(\Psi_{1,0}\frac{k_m}{\gamma_m} + \Psi_{1,1}F_+^{(1)}(1)\right) = \frac{\frac{k_m}{\gamma_m}\frac{1}{2}\Psi_{1,0}}{1 - \frac{1}{2}\Psi_{1,1}} \tag{36}$$

Proceeding similarly we have $F_+^{(2)}(1)$ determined by $F_+^{(1)}(1)$ as follows:

$$F_+^{(2)}(1) = \frac{\left(\frac{k_m}{\gamma_m}\right)^2 \frac{1}{2^2}\Psi_{2,0}}{1 - \frac{1}{2^2}\Psi_{2,2}} + \frac{\left(\frac{k_m}{\gamma_m}\right)^2 \frac{1}{2^3}\binom{2}{1}\Psi_{2,1}\Psi_{1,0}}{(1-\frac{1}{2}\Psi_{1,1})(1-\frac{1}{2^2}\Psi_{2,2})} \tag{37}$$

Next, the coefficient

$$\begin{aligned}
F^3(1) = &\left[\frac{\frac{1}{2^3}\left(\frac{k_m}{\gamma_m}\right)^3 \Psi_{3,0}}{1-\frac{1}{2^3}\Psi_{3,3}}\right] + \left[\frac{\frac{1}{2^4}\binom{3}{1}\Psi_{3,1}\Psi_{1,0}\left(\frac{k_m}{\gamma_m}\right)^3}{(1-\frac{1}{2^3}\Psi_{3,3})(1-\frac{1}{2}\Psi_{1,1})}\right] + \left[\frac{\frac{1}{2^5}\binom{3}{2}\Psi_{3,2}\Psi_{2,0}\left(\frac{k_m}{\gamma_m}\right)^3}{(1-\frac{1}{2^2}\Psi_{2,2})(1-\frac{1}{2^3}\Psi_{3,3})}\right] \\
&+ \left[\frac{\frac{1}{2^6}\binom{3}{2}\Psi_{3,2}\Psi_{2,1}\Psi_{1,0}\left(\frac{k_m}{\gamma_m}\right)^3}{(1-\frac{1}{2}\Psi_{1,1})(1-(\frac{1}{2^2}\Psi_{2,2})(1-\frac{1}{2^3}\Psi_{3,3})}\right]
\end{aligned} \tag{38}$$

Observing the pattern of the successive coefficients, we obtain the general solution for $F_+^{(k)}(1)$ as follows:

$$F_+^{(k)}(1) = \left(\frac{k_m}{\gamma_m}\right)^k \frac{1}{2^k}\frac{1}{(1-\frac{1}{2^k}\Psi_{k,k})}\left[\sum_{\{S_{k-1}\}} \frac{(\frac{1}{2})^{\sum_i j_i}\phi_{k,j_z}\ \phi_{j_z,j_{z-1}} \dots \phi_{j_{1,0}}}{\prod_i(1-\frac{1}{2^{j_i}}\Psi_{j_i,j_i})} + \phi_{k,0}\right] \tag{39}$$

where $\phi_{k,j} = \Psi_{k,j}\binom{k}{j}$. Here $\{S_{k-1}\}$ denotes the set of all the subsets $S_{k-1} = \{j_i\} = (j_z, j_{z-1}...j_1)$ of integers $j_i \in (1, 2, ...k-1)$ such that $j_z > j_{z-1} > ... > j_1$. For example for $k = 3$, the subsets are (1), (2), and (2,1) as is seen in Eq. 38.

With the coefficients given by Eq. 39, the cyclo-stationary distributions are formally given by the series:

$$P_+^{ss}(m_+) = \sum_{k=m_+}^{\infty} \binom{k}{m_+} \frac{(-1)^{k-m_+}}{k!} F_+^{(k)}(1) \tag{40}$$

$$P_-^{ss}(m_-) = \sum_{k=m_-}^{\infty} \binom{k}{m_-} \frac{(-1)^{k-m_-} 2^k}{k!} F_+^{(k)}(1) \tag{41}$$

##### C. The cyclo-stationary mRNA distributions are Poisson for fixed cell-division times $T$

For fixed cell division times, $g(t_s) = \delta(t_s - T)$, we have $\Psi_{k,j} = e^{-j\gamma_m T}(1 - e^{-\gamma_m T})^{k-j}$, and

$$\begin{aligned}\Psi_{k,j_z}\Psi_{j_z,j_{z-1}}\dots\Psi_{j_1,0} &= e^{-j_z\gamma_m T}(1 - e^{-\gamma_m T})^{k-j_z} e^{-j_{z-1}\gamma_m T}(1 - e^{-\gamma_m T})^{j_z - j_{z-1}} \dots . e^{-0*\gamma_m T}(1 - e^{-\gamma_m T})^{j_1} \\ &= e^{-\gamma_m T \sum_{i=1}^{z} j_i}(1 - e^{-\gamma_m T})^k \end{aligned} \tag{42}$$

Consequently from Eqs. 40 and 39,

$$\begin{aligned} P_+^{ss}(m_+) &= \sum_{k=m_+}^{\infty} \frac{(-1)^{k-m_+}}{k!} \binom{k}{m_+} F_+^{(k)}(1) \\ &= \sum_{k=m_+}^{\infty} \frac{(-1)^{j-m_+}}{k!} \binom{k}{m_+} \left[ \left(\frac{k_m}{\gamma_m}\right)^k \frac{1}{2^k} \frac{1}{(1 - \frac{1}{2^k}\Psi_{k,k})} \left[ \sum_{\{S_{k-1}\}} \frac{(\frac{1}{2})^{\sum j_i} \phi_{k,j_z}\phi_{j_z,j_{z-1}}...\phi_{j_1,0}}{\prod_i (1 - \frac{1}{2^{j_i}}\Psi_{j_i,j_i})} + \phi_{k,0} \right] \right] \\ &= \sum_{k=m_+}^{\infty} \frac{(-1)^{k-m_+}}{j!} \binom{k}{m_+} \left(\frac{k_m}{\gamma_m}\right)^k \frac{\frac{1}{2^k}}{1 - \frac{1}{2^k}\Psi_{k,k}} (1 - e^{-\gamma_m T})^k \left[ 1 + \sum_{\{S_{k-1}\}} \binom{k}{j_z}\binom{j_z}{j_{z-1}} ... \binom{j_2}{j_1} \frac{\prod_i (\frac{1}{2}e^{-\gamma_m T})^{j_i}}{\prod_i (1 - (\frac{1}{2}e^{-\gamma_m T})^{j_i})} \right] \end{aligned} \tag{43}$$

Using the the following identity [1], with $x = \frac{1}{2}e^{-\gamma_m T}$ in this case,

$$1 + \sum_{\{S_{k-1}\}} \binom{k}{j_z}\binom{j_z}{j_{z-1}} ... \binom{j_2}{j_1} \prod_i \frac{x^{j_i}}{1 - x^{j_i}} = \frac{1 - x^k}{(1 - x)^k} \tag{44}$$

Thus the $(1 - x^k)$ factors cancel from the numerator and denominator, and Eq. 43 simplifies to

$$\begin{aligned} P_{ss}(m_+) &= \sum_{k=m_+}^{\infty} \binom{k}{m_+} \frac{(-1)^{k-m_+}}{k!} \left(\frac{k_m}{\gamma_m}\right)^k \frac{\frac{1}{2^k}}{(1 - e^{-\gamma_m T}\frac{1}{2})^k} (1 - e^{-\gamma_m T})^k \\ &= \sum_{k=m_+}^{\infty} \frac{d^k}{m_+!(k - m_+)!} (-1)^{k-m_+} \end{aligned} \tag{45}$$

with $d = \left(\frac{k_m}{\gamma_m}\right) \frac{(1-e^{-\gamma_m T})}{(2-e^{-\gamma_m T})}$. Thus $F_+^{(k)}(1) = d^k$. Replacing $k - m_+ = r$, then Eq. 45 reduces to a Poisson distribution:

$$P_+^{ss}(m_+) = \frac{d^{m_+}}{m_+!} \sum_{r=0}^{\infty} \frac{(-1)^r d^r}{r!} = \frac{d^{m_+}}{m_+!} \exp(-d). \tag{46}$$

Since Eq. 41 has an extra factor of $2^k$ multiplying $F_+^{(k)}(1)$, we would have $d$ replaced by $2d$ and the distribution:

$$P_-^{ss}(m_-) = \frac{(2d)^{m_-}}{m_-!} \exp(-2d). \tag{47}$$

##### D. Simplified form of the cyclo-stationary distributions for Exponentially distributed cell cycle times

For exponentially distributed division times, $g(t_s) = \lambda e^{-\lambda t_s}$ (with $\lambda = 1/T$), and

$$\Psi_{k,j} = \int_0^\infty dt_s g(t_s) e^{-\gamma_m t_s}(1-e^{-\gamma_m t_s})^{k-j} = \lambda/\gamma_m\, B\left(\frac{\lambda + j\gamma_m}{\gamma_m}, k-j+1\right) = \frac{\lambda}{\gamma_m}\frac{\Gamma(k-j+1)\Gamma\left(\frac{\lambda}{\gamma_m}+j\right)}{\Gamma\left(\frac{\lambda}{\gamma_m}+k+1\right)} \tag{48}$$

and $\Psi_{k,k} = \frac{\lambda}{\gamma_m}\frac{\Gamma(1)\Gamma\left(\frac{\lambda}{\gamma_m}+k\right)}{\Gamma\left(\frac{\lambda}{\gamma_m}+k+1\right)} = \frac{\lambda}{\lambda+\gamma_m k}$. Then

$$\phi_{k,j} = \binom{k}{j}\Psi_{k,j} = \left(\frac{\lambda}{\gamma_m}\right)\frac{\Gamma(k+1)\Gamma[\frac{\lambda}{\gamma_m}+j]}{\Gamma(j+1)\Gamma[\frac{\lambda}{\gamma_m}+k+1]} \tag{49}$$

Hence $\phi_{k,0} = k!\frac{\Gamma\left(\frac{\lambda}{\gamma_m}\right)}{\Gamma\left(\frac{\lambda}{\gamma_m}+k+1\right)}$. Furthermore,

$$\phi_{n,j_z}\ \phi_{j_z,j_{z-1}}\ \ldots\ \phi_{j_1,0} = \frac{\lambda^{z+1}}{\gamma_m^{z+1}}\frac{k!\ \Gamma[\frac{\lambda}{\gamma_m}]}{\Gamma[\frac{\lambda}{\gamma_m}+k+1]}\prod_{i=1}\frac{\gamma_m}{\lambda+\gamma_m j_i} \tag{50}$$

Substituting the above, in Eq. 40 and 39, we have

$$\begin{aligned}
P_+^{ss}(m_+) &= \sum_{k=m_+}^{\infty}\binom{k}{m_+}(-1)^{k-m_+}\frac{1}{k!}\left(\frac{k_m}{\gamma_m}\right)^k\frac{1/2^k}{(1-1/2^k\Psi_{k,k})}\left[\sum_{\{S_{k-1}\}}\frac{(\frac{1}{2})^{\sum j_i}\phi_{k,j_z}\ \phi_{j_z,j_{z-1}}\ \ldots\ \phi_{j_1,0}}{\prod_i(1-\frac{1}{2^{j_i}}\Psi_{j_i,j_i})}+\phi_{k,0}\right]\\
&= \sum_{k=m_+}^{\infty}\binom{k}{m_+}(-1)^{k-m_+}\frac{\frac{1}{2^k}\left(\frac{k_m}{\gamma_m}\right)^k}{(1-\frac{1}{2^k}\Psi_{k,k})}\left[\frac{\lambda}{\gamma_m}\frac{\Gamma(\frac{\lambda}{\gamma_m})}{\Gamma(\frac{\lambda}{\gamma_m}+k+1)}+\sum_{\{S_{k-1}\}}\frac{(\frac{1}{2})^{\sum j_i}\frac{\lambda^{z+1}}{\gamma_m^{z+1}}\frac{\Gamma(\lambda/\gamma_m)}{\Gamma(\lambda/\gamma_m+k+1)}}{\prod_i(1-\frac{1}{2^{j_i}}\Psi_{j_i,j_i})}\prod_i\frac{\gamma_m}{\lambda+\gamma_m j_i}\right]\\
&= \sum_{k=m_+}^{\infty}\binom{k}{m_+}(-1)^{k-m_+}\frac{\frac{1}{2^k}\left(\frac{k_m}{\gamma_m}\right)^k}{(1-\frac{1}{2^k}\Psi_{k,k})}\frac{\Gamma(\frac{\lambda}{\gamma_m}+1)}{\Gamma(\frac{\lambda}{\gamma_m}+k+1)}\left[1+\sum_{\{S_{k-1}\}}\prod_i\frac{\frac{1}{2^{j_i}}\Psi_{j_i,j_i}}{(1-\frac{1}{2^{j_i}}\Psi_{j_i,j_i})}\right]\\
&= \sum_{k=m_+}^{\infty}\binom{k}{m_+}(-1)^{k-m_+}\frac{\frac{1}{2^k}\left(\frac{k_m}{\gamma_m}\right)^k}{(1-\frac{1}{2^k}\Psi_{k,k})}\frac{\Gamma(\frac{\lambda}{\gamma_m}+1)}{\Gamma(\frac{\lambda}{\gamma_m}+k+1)}\frac{1}{\prod_i^{k-1}(1-\frac{1}{2^i}\Psi_{i,i})}\\
&= \sum_{k=m_+}^{\infty}\binom{k}{m_+}\frac{(-1)^{k-m_+}}{k!}\frac{k!\left(\frac{k_m}{2}\right)^k}{\prod_i^k(\lambda+\gamma_m i-\frac{1}{2^i}\lambda)},
\end{aligned} \tag{51}$$

where we have used an identity [1], with $f(j_i) = \frac{1}{2^{j_i}}\Psi_{j_i,j_i}$ as follows:

$$1+\sum_{\{S_{k-1}\}}\prod_i\frac{f(j_i)}{(1-f(j_i))} = \frac{1}{\prod_i^{k-1}(1-f(i))} \tag{52}$$

Eq. 51 shows that $F_+^{(k)}(1) = k!\left(\frac{k_m}{2}\right)^k/\prod_i^k(\lambda+\gamma_m i-\frac{1}{2^i}\lambda)$. For obtaining the distribution before division, we note the extra factor of $2^k$ in Eq. 41, and that implies (comparing with Eq. 51)

$$P_-^{ss}(m_-) = \sum_{k=m_-}^{\infty}\binom{k}{m_-}\frac{(-1)^{k-m_-}}{k!}\frac{k!\,(k_m)^k}{\prod_i^k(\lambda+\gamma_m i-\frac{1}{2^i}\lambda)}. \tag{53}$$

##### E. Expressions of $CV^2$ and Skewness of the distribution $P_+^{ss}(m_+)$ of mRNA at cell birth.

Using the exact expressions of $F_+^{(1)}(1)$, $F_+^{(2)}(1)$, and $F_+^{(3)}(1)$ in Eqs. 36, 37 and 38, we have obtained the cumulants (from Eqs. 24, 26 and 28), and thus studied the mean, $CV^2$ and Skewness in the main text.

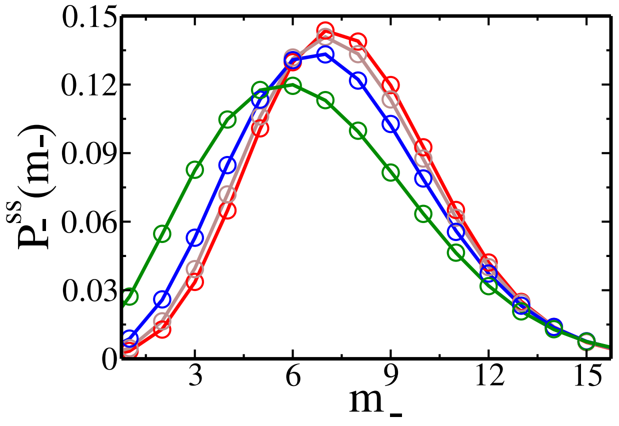


FIG. 1. Cyclo-stationary distribution $P_{-}^{ss}(m_{-})$ of mRNA for the four $g(t_s)$ shown in Fig. 3 in the main text (corresponding colours being the same)

#### III. STATISTICS OF CYCLO-STATIONARY PROTEIN COUNT

##### A. The generating function for protein kinetics, and thereby obtaining function $\mathcal{H}$

The Master equation for the bursty protein translation is the following (with initial copy number being $n'_{+}$):

$$\frac{\partial P(n,t|n'_{+})}{\partial t} = k_m \left[\sum_{r=1}^{n} \frac{b^r}{(b+1)^{r+1}} P(n-r,t|n'_{+}) - \frac{b}{b+1} P(n,t|n'_{+})\right] + \gamma_p \left[(n+1)P(n+1,t|n'_{+}) - nP(n,t|n'_{+})\right]. \tag{54}$$

The generating function $F(q,t|n'_{+}) = \sum_{j=0}^{\infty} q^n p(n,t|n'_{+})$ is known to satisfy [2]

$$\frac{1}{v}\frac{\partial F}{\partial \tau} + \frac{\partial F}{\partial v} = \frac{ab}{1-bv} F \tag{55}$$

where $v = q-1$, $\tau = \gamma_p t$ and $a = \frac{k_m}{\gamma_p}$. The solution of Eq. 54 by the method of Lagrange characteristics yield [2]

$$F(q,t|n'_{+}) = \left(\frac{1-b(q-1)e^{-\gamma_p t_s}}{1-b(q-1)}\right)^a \times \left(1+(q-1)\,e^{-\gamma_p t_s}\right)^{n'_{+}}. \tag{56}$$

Thus comparing with Eq. 5 of the main text (also see below Eq. 9), we identify the function

$$\mathcal{H} = \left(\frac{1-b(q-1)e^{-\gamma_p t_s}}{1-b(q-1)}\right)^a. \tag{57}$$

##### B. Obtaining the recursion relation for the coefficients $F_+^{(k)}(1)$ in the series of the distributions $P_\pm^{ss}(m_\pm)$

Using $\mathcal{H}$ from Eq. 57 in Eq. 10, and $F_+(q) = \sum_{j=0}^{\infty} \frac{(q-1)^j}{j!} F_+^{(j)}(1)$ we have

$$\begin{aligned}
F_+(q) &= \int_0^\infty dt_s g(t_s) \left( \frac{1 - \frac{b}{2}(q-1)e^{-\gamma_p t_s}}{1 - \frac{b}{2}(q-1)} \right)^a F_+\left(1 + \frac{(q-1)}{2} e^{-\gamma_p t_s}\right) \\
&= \int_0^\infty dt_s g(t_s) \left( \frac{1 - \frac{b}{2}(q-1)e^{-\gamma_p t_s}}{1 - \frac{b}{2}(q-1)} \right)^a \sum_{j=0}^{\infty} \frac{F_+^j(1)\ (q-1)^j}{2^j j!} e^{-j\gamma_p t_s} \\
&= \int_0^\infty dt_s g(t_s) \sum_{l=0}^{\infty} \frac{\Gamma(a+1)}{\Gamma(a-l+1)} \left(\frac{b}{2}\right)^l \frac{(-1)^l (q-1)^l e^{-l\gamma_p t_s}}{l!} \sum_{s=0}^{\infty} \frac{\Gamma(a+s)}{\Gamma(a)} \left(\frac{b}{2}\right)^s \frac{(q-1)^s}{s!} \sum_{j=0}^{\infty} \frac{F_+^j(1) e^{-j\gamma_p t_s} (q-1)^j}{2^j j!} \\
&= \sum_{l=0}^{\infty}\sum_{j=0}^{\infty}\sum_{s=0}^{\infty} \int_0^\infty dt_s g(t_s) (-1)^l \frac{\Gamma(a+1)}{\Gamma(a-l+1)} \frac{\Gamma(a+s)}{\Gamma(a)} \left(\frac{b}{2}\right)^{l+s} (q-1)^{l+j+s} e^{-(l+j)\gamma_p t_s} \binom{a+k-1}{k} \frac{F_+^{(j)}(1)}{2^j\ l!\ s!\ j!}
\end{aligned} \tag{58}$$

We define $L_{l+j} = \int_0^\infty dt_s g(t_s) e^{-(l+j)\gamma_p t_s}$, and replace indices $l+j+s=k$, which leads to

$$F_+(q) = \sum_{k=0}^{\infty} \frac{(q-1)^k}{k!} \sum_{l=0}^{k} \sum_{j=0}^{k-l} a \frac{k!\Gamma(a+k-l-j)}{\Gamma(a-l+1)} (-1)^l \left(\frac{b}{2}\right)^{k-j} \frac{L_{l+j} F_+^{(j)}(1)}{l! j! (k-l-j)! 2^j} \tag{59}$$

A series expansion on the left side of Eq. 59 about $q=1$, and comparing with the right side, yields the desired recursion relation:

$$F_+^{(k)}(1) = ak! \sum_{l=0}^{k} \sum_{j=0}^{k-l} (-1)^l \frac{b^{k-j}}{2^k} L_{l+j} \frac{(a+k-l-j-1)!}{(a-l)!\ l!\ j!\ (k-l-j)!} F_+^{(j)}(1) \tag{60}$$

The above Eq. 60 appears as Eq. 26 in the main text. Once these coefficients $F_+^{(k)}(1)$ are solved for, they may be used to obtain the cyclo-stationary distributions

$$P_+^{(ss)}(n_+) = \sum_{k=n_+}^{\infty} \binom{k}{n_+} \frac{(-1)^{k-n_+}}{k!} F_+^{(k)}(1) \tag{61}$$

$$P_-^{(ss)}(n_-) = \sum_{k=n_-}^{\infty} \binom{k}{n_-} \frac{(-1)^{k-n_-} 2^k}{k!} F_+^{(k)}(1) \tag{62}$$

##### C. Expressions of first three $F_+^{(k)}(1)$ which determine exactly $CV^2$ and Skewness of the distribution $P_+^{ss}(n_+)$ of protein.

Firstly, $F_+^{(0)}(1) = 1$. Then using the above Eq. 60 recursively, we get

$$F_+^{(1)}(1) = ab \frac{1-L_1}{2-L_1} \tag{63}$$

then,

$$F_+^{(2)}(1) = \frac{ab^2}{4-L_2} \left[ (1-L_2) + a \frac{(2-3L_1+L_1L_2)}{(2-L_1)} \right] \tag{64}$$

and then,

$$\begin{aligned}
F_+^{(3)}(1) = \frac{1}{8-L_3} \Big[ & ab^3 \Big( (1+a)(2+a) - 3a((1+a)L_1 - (a-1)L_2) - (a-2)(a-1)L_3 \Big) \\
& + 3ab^2 \Big( (1+a)L_1 - 2aL_2 + (a-1)L_3 \Big) F_+^{(1)}(1) + 3ab \Big( L_2 - L_3 \Big) F_+^{(2)}(1) \Big]
\end{aligned} \tag{65}$$

The constants $L_1$, $L_2$ and $L_3$ may be evaluated given a $g(t_s)$. Then the above Eqs. 63, 64 and 65 are substituted in the Eqs. 24, 26 and 28, to obtain the cumulants and thus $CV^2$ and Skewness, which are studied in the main text.

#### IV. COMPUTATIONAL METHODS

##### A. Precautions to perform numerical sums of different series to obtain the coefficients $F_+^{(k)}(1)$ and the theoretical cyclo-stationary distributions $P_\pm^{ss}$

In this work we had to sum various series to obtain the desired coefficients and functions.

The equations for the coefficients $F_+^{(k)}(1)$ appear as Eq. 35 for mRNA, and Eq. 60 for protein, and are of the form

$$F_+^{(k)}(1) = \sum_{j=1}^{k-1} c_{k,j} F_+^{(j)}(1) \tag{66}$$

As the values of $F_+^{(j)}(1)$ grow very fast with $j$ we loose precision soon in ordinary calculations. A better way to store large numbers is by taking logarithm, and we do so for terms in Eq. 66. Thus we store terms

$$u_{k,j} = \ln c_{k,j} + \ln F_+^{(j)}(1). \tag{67}$$

We specify very high precision for such calculation and storage in Mathematica (through the SetPrecision[d] command) up to $d = 100$ decimal places in case of mRNA and $d = 200$ decimal places for protein. We reconstruct back the coefficient

$$F_+^{(k)}(1) = \sum_{j=1}^{k-1} e^{u_{k,j}}. \tag{68}$$

Once the coefficients $F_+^{(k)}(1)$ are obtained by the above method, up to some desired $k$, we put them in the series in Eqs. 40 and 41 for mRNA, and Eqs. 61, 62 for protein, to obatin the cyclo-stationary distributions. For mRNA, convergence was attained for $\sim 30-50$ terms in the series of $P_\pm^{ss}(m_\pm)$.

For protein, ordinary sum of the series of $P_\pm^{ss}(n_\pm)$ were not enough with reasonable values of $k$. We used Borel sum formula as follows:

$$P_\pm^{ss}(n_\pm) = \sum_{k=n_\pm}^{\infty} f(k, n_\pm) = P_\pm^{ss}(n_\pm)\Big|_{\text{Borel}} = \lim_{t\to\infty} e^{-t} \sum_{n=0}^{M\to\infty} \frac{t^n}{n!} \sum_{k'=0}^{n} f(k'+n_\pm, n_\pm) \tag{69}$$

In calculations we choose $M \sim 200-250$ and $t = 30$ to obtain convergence of the protein cyclo-stationary distributions.

##### B. Kinetic Monte Carlo Simulations

We perform Kinetic Monte Carlo (KMC) or Gillespie [3] simulations for the various models governing the transcription or translation models of mRNA and protein this paper, undergoing Binomial partitioning after random time intervals $t_s$ drawn from some distribution $g(t_s)$. Thus, at any instant, there are three possible events to either increase, decrease, or reset the copy number (due to cell partition). We typically use $\sim 10^7$ histories for getting the data for various distributions and cumulants, which were then compared with the theory.

#### V. AGE DEPENDENT CYCLO-STATIONARY DISTRIBUTIONS

The cyclo-stationary distribution $P^{ss}(y, \tau)$ of cells at an age $\tau$ before the next cell division, may written with respect to $P_+^{ss}(y_+)$ at birth, as follows:

$$P^{ss}(y, \tau) = \sum_{y_+} P_+^{ss}(y_+) p(y, \tau | y_+). \tag{70}$$

Using its generating function $\tilde{G}(q,\tau) = \sum_y q^y P^{ss}(y,\tau)$, from Eq. 70 (and using the same steps as in Eqs. 9 and 10)

$$\tilde{G}(q,\tau) = \sum_{y_+} P_+^{ss}(y_+) F(q,\tau|y_+) = \mathcal{H}(q-1,\gamma_y\tau)\ F_+(1+(q-1)e^{-\gamma_y\tau})$$
$$= \sum_k \frac{(q-1)^k}{k!} G_y^{(k)}(\tau) \tag{71}$$

The defined quantities $G_y^{(k)}(\tau)$ are obtained in Eq. 71 by expressing $F_+(q) = \sum_{j=0}^{\infty} \frac{(q-1)^j}{j!} F_+^{(j)}(1)$, and expanding $\mathcal{H}$ as a power series of $(q-1)$. The resulting expression of $G_y^{(k)}(\tau)$ are of the form of the integrands of Eqs. 33 or 58 (without the $\int_0^\infty dt_S g(t_s)/2^k$ factors), and are explicitly given for mRNA and protein in Eqs. 30 and 31 in the main text. Finally it is easy to obtain the desired age-dependent distributions in terms of $G_y^{(k)}(\tau)$ as

$$P^{ss}(y,\tau) = \frac{1}{y!}\frac{\partial^y \tilde{G}}{\partial q^y}\bigg|_{q=0} = \sum_{k=y}^{\infty} \binom{k}{y} \frac{(-1)^{k-y}}{k!} G_y^{(k)}(\tau). \tag{72}$$

#### VI. GENERATING FUNCTIONS OF PROTEIN DISTRIBUTION AT BIRTH, FOR DETERMINISTIC PARTITIONING, AND DETERMINISTIC GENE EXPRESSION

If we have a deterministic partitioning, we would replace the binomial distribution $B(\tilde{y}_+, \frac{1}{2}, x_+)$ in Eqs. 8 and 9 by $\delta_{x_+,\tilde{y}_+/2}$ and as a result

$$F_+(q) = \sum_{y'_+} \int_0^\infty dt_s g(t_s) \sum_{\tilde{y}_+} q^{\tilde{y}_+} p(\tilde{y}_+, t_s|y'_+) \sum_{x_+=0}^{\tilde{y}_+} \frac{1}{q^{x_+}} \delta_{x_+,\frac{\tilde{y}_+}{2}} P_+^{ss}(y'_+)$$
$$= \sum_{y'_+} \int_0^\infty dt_s g(t_s) \left( \sum_{\tilde{y}_+} q^{\tilde{y}_+/2} p(\tilde{y}_+, t_s|y'_+) \right) P_+^{ss}(y'_+)$$
$$= \int_0^\infty dt_s g(t_s) \sum_{y'_+} F\left(\sqrt{q}, t_s \middle| y'_+\right) P_+^{ss}(y'_+). \tag{73}$$

For proteins $y \equiv n$, and using the appropriate $F(q,t|n'_+)$ from Eq. 56, we have the counterpart of Eq. 10 as:

$$F_+(q) = \int_0^\infty dt_s g(t_s) \left( \frac{1-b\left(\sqrt{q}-1\right)e^{-\gamma_p t_s}}{1-b\left(\sqrt{q}-1\right)} \right)^a F_+\left(1+(\sqrt{q}-1)e^{-\gamma_p t_s}\right) \tag{74}$$

If in addition to deterministic partitioning, one also has deterministic protein kinetics

$$\frac{dn}{dt} = k_m b - \gamma_p n, \tag{75}$$

then $\tilde{n}_+ = n'_+ e^{-\gamma_p t_s} + ab(1-e^{-\gamma_p t_s})$ and $F$ in Eq. 73 gets replaced by $q^{\frac{1}{2}(\lambda_p + n'_+ e^{-\gamma_p t_s})}$ where $\lambda_p = ab(1-e^{-\gamma_p t_s})$, i.e.

$$F_+(q) = \sum_{n'_+} \int_0^\infty dt_s g(t_s) \left( \sum_{\tilde{n}_+} q^{\tilde{n}_+/2} p(\tilde{n}_+, t_s|n'_+) \right) P_+^{ss}(n'_+)$$
$$= \int_0^\infty dt_s g(t_s) \sum_{n'_+} P_+^{ss}(n'_+) q^{\frac{1}{2}(\lambda_p + n'_+ e^{-\gamma_p t_s})}$$
$$= \int_0^\infty dt_s g(t_s) q^{\frac{1}{2}\lambda_p} \sum_{n'_+} P_+^{ss}(n'_+) \left(q^{\frac{1}{2}e^{-\gamma_p t_s}}\right)^{n'_+}$$
$$= \int_0^\infty dt_s g(t_s) q^{\frac{1}{2}\lambda_p} F_+(q^{\frac{1}{2}e^{-\gamma_p t_s}}) \tag{76}$$

The moment $\langle n_+ \rangle = q\frac{\partial}{\partial q}F_+(q)\Big|_{q=1}$ and $\langle n_+^2 \rangle = q\frac{\partial}{\partial q}q\frac{\partial}{\partial q}F_+(q)\Big|_{q=1}$, and hence taking derivatives of on two sides of Eq. 74 and 76 respectively, and setting $q = 1$, we may obtain the moments in the two cases above. The explicit forms of $CV^2$ thus obtained are shown in Eq. 32 in the main text (corresponding to Eq. 74) and in Eq. 34 in the main text (corresponding to Eq. 76).

[1] K. Rijal, N. I. C. Müller, E. Friauf, A. Singh, A. Prasad, and D. Das, Phys. Rev. Lett. **132**, 228401 (2024).
[2] V. Shahrezaei and P. S. Swain, Proc. Natl. Acad. Sci. USA **105**, 17256 (2008).
[3] D. T. Gillespie, J. Phys. Chem. **81**, 2340 (1977).
